# Tumor-resident regulatory T cells in pancreatic cancer express the αvβ5 integrin as a targetable activation marker

**DOI:** 10.1101/2023.05.24.542137

**Authors:** Kodai Suzuki, Yuki Kunisada, Norio Miyamura, Shingo Eikawa, Tatiana Hurtado de Mendoza, Evangeline S. Mose, Caisheng Lu, Yukihito Kuroda, Erkki Ruoslahti, Andrew M. Lowy, Kazuki N. Sugahara

## Abstract

Pancreatic ductal adenocarcinoma (PDAC) has abundant immunosuppressive regulatory T cells (Tregs), which contribute to a microenvironment resistant to immunotherapy. Here, we report that Tregs in the PDAC tissue, but not those in the spleen, express the αvβ5 integrin in addition to neuropilin-1 (NRP-1), which makes them susceptible to the iRGD tumor-penetrating peptide, which targets cells positive for αv integrin- and NRP-1. As a result, long-term treatment of PDAC mice with iRGD leads to tumor-specific depletion of Tregs and improved efficacy of immune checkpoint blockade. αvβ5 integrin^+^ Tregs are induced from both naïve CD4^+^ T cells and natural Tregs upon T cell receptor stimulation, and represent a highly immunosuppressive subpopulation of CCR8^+^ Tregs. This study identifies the αvβ5 integrin as a marker for activated tumor-resident Tregs, which can be targeted to achieve tumor-specific Treg depletion and thereby augment anti-tumor immunity for PDAC therapy.

## INTRODUCTION

Pancreatic ductal adenocarcinoma (PDAC) is poorly responsive to conventional immune checkpoint therapy, at least in part, due to the prevalence of immunosuppressive cells and paucity of cytotoxic effector cells^1–3^. Regulatory T cells (Tregs) are believed to be a significant driver of the immunosuppressive microenvironment in the PDAC tissue that protects the cancer cells from T cell mediated anti-tumor immunity^4^. In fact, a decreased ratio of tumor-infiltrating CD8^+^ T cells to Tregs (CD8/Treg ratio) correlates with poor prognosis in PDAC patients^5^ and diminished response to immune checkpoint blockade (ICB) therapy^6^.

Various therapeutic approaches have been taken to deplete tumor-resident Tregs. Examples include the use of inhibitors of cytokines and chemokines that recruit Tregs to PDAC tissue, antibodies (Abs) that target cell surface molecules expressed on Tregs such as CD25, cytotoxic T-lymphocyte-associated protein 4 (CTLA-4), programmed cell death-1 (PD-1), and the glucocorticoid induced TNF receptor^7–10^. Small molecules that kill highly proliferating cells have also been used^11, 12^. However, most of these approaches lead to systemic Treg depletion, which can result in serious autoimmune related toxicities^13, 14^. A study using transgenic *Kra*s mutant mice showed that systemic Treg depletion unexpectedly accelerated tumor progression by inducing tumor infiltration of immunosuppressive myeloid cells^15^. These findings suggest that Treg depletion must be achieved in a controlled, ideally tumor-specific, manner to avoid autoimmune complications and unwanted alterations of the tumor immune microenvironment.

The benefits of tumor-specific Treg depletion in cancer therapy have been demonstrated by targeting the chemokine receptor CCR8^16, 17^. CCR8 is a recently described marker for tumor-resident Tregs, which is expressed on up to 80% of tumor-resident Tregs but minimally on other immune cells^16, 18, 19^. CCR8 appears to be expressed on Tregs in response to antigen recognition, and reflects enhanced immunosuppressive functions of these Tregs^16, 18–20^. Depleting CCR8^+^ Tregs leads to improved anti-tumor immunity without causing autoimmune complications, reinforcing the importance of tumor selectivity when Treg depletion is used in cancer treatment^16^.

iRGD is a cyclic tumor-penetrating peptide (amino acid sequence: CRGDKGPDC) that delivers drugs deep into the extravascular tumor tissue in a tumor-specific manner^21^. It carries a tumor-specific RGD motif that binds to αvβ3/β5 integrins and a tissue/cell-penetrating RXXK/R motif that binds to neuropilin-1 (NRP-1). Systemically injected iRGD homes to tumors by targeting the αvβ3/β5 integrins expressed on tumor endothelial cells. It is then proteolytically processed to expose an active RXXK/R motif that now binds to NRP-1. The RXXK/R-NRP-1 interaction activates a transcytotic penetration pathway in the tumor, which is mediated by vesicles that resemble macropinosomes^22, 23^. This mechanism allows iRGD as well as co-injected bystander molecules to penetrate through cell layers and widely spread into the tumor tissue^24^.

Our recent work revealed that the expression of αvβ5 integrin in the tumor (in addition to NRP-1) is critical for iRGD to effectively penetrate extravascular tumor tissue^25^. Previous studies have shown that the αvβ5 integrin is expressed in ∼80% of human PDAC tissues and is a predictor of poor prognosis in PDAC patients^26^. In fact, epithelial cells and cancer-associated fibroblasts (CAFs) in the PDAC tissue often express high levels of αvβ5 integrin and NRP-1, and their crosstalk mediated by transforming growth factor-β (TGF-β) helps maintain the expression of αvβ5 integrin, creating a tumor microenvironment optimal for iRGD penetration^25^. Therefore, iRGD co-injection therapy is highly effective in allowing for tumor drug penetration in PDAC despite the desmoplastic nature of the tumor. In a recent phase 1b clinical trial, iRGD (in the name of CEND-1) showed promising preliminary efficacy in combination with gemcitabine (Gem) and nab-paclitaxel (Nab-P) against metastatic PDAC^27^. iRGD is now being evaluated in multiple phase 2 clinical trials for PDAC and other cancers (e.g., NCT05042128).

In this article, we report that the αvβ5 integrin is also expressed on Tregs that infiltrate the PDAC tissue, but not those in the spleen, allowing iRGD to selectively target PDAC-resident Tregs. Importantly, the αvβ5 integrin defines a highly immunosuppressive subpopulation of CCR8^+^ Tregs, providing a new way to target activated Tregs in a tumor-specific manner.

## RESULTS

### iRGD monotherapy reduces Tregs in the PDAC tissue but not in the spleen

We have previously reported that iRGD + Gem significantly prolonged the survival of transgenic *Kras^G12D/+^;LSL-Trp53^R172H/+^;Pdx-1-Cre* (KPC) PDAC mice compared to iRGD or Gem therapy alone^25^. Further analysis revealed that the samples from the iRGD + Gem arm harbored significantly more CD8^+^ T cells than those from the Gem monotherapy arm (Supplementary Fig. 1). Although less effective than iRGD + Gem, iRGD monotherapy also increased the number of CD8^+^ T cells in the PDAC tissue, suggesting that iRGD has an immunomodulatory effect. We hypothesized that the effect was a response to Treg targeting enabled by iRGD given that mouse Tregs express NRP-1^28^, one of the iRGD receptors^21^. The hypothesis appeared reasonable because there were less PDAC-resident cells expressing Foxp3, a master regulator of Tregs^29, 30^, in the iRGD monotherapy and iRGD + Gem combination therapy arms compared to the others (Supplementary Fig. 2).

To study the hypothesis further, we generated a syngeneic PDAC mouse model using luciferase^+^ PDAC cells derived from KPC mice. The PDAC tumors rapidly grew orthotopically in B6129SF1/J hybrid mice (Supplementary Fig. 3). In line with previous studies^21, 24, 31^, iRGD monotherapy did not affect tumor growth (Supplementary Fig. 4). However, flow cytometry (Supplementary Fig. 5) revealed that 7 days of iRGD treatment significantly reduced the proportion of Tregs (Fig. 1a, left panel; Supplementary Fig. 6, top panels), and increased the proportion of CD8^+^ T cells (Supplementary Fig. 7a) and the CD8/Treg ratio in the PDAC (Fig. 1a, right panel). Similar results were obtained by immunofluorescence (IF) (Fig. 1b). In contrast, iRGD monotherapy did not affect the proportion of Tregs or CD8^+^ T cells or the CD8/Treg ratio in the spleen of the PDAC mice (Fig. 1c; Supplementary Fig. 6, bottom panels; Supplementary Fig. 7b). The data suggest that the effect of iRGD on the immune cells was tumor specific. At baseline, the CD8/Treg ratio was decreased in the spleen when the mice carried a PDAC (Supplementary Fig. 8).

**Figure 1.**
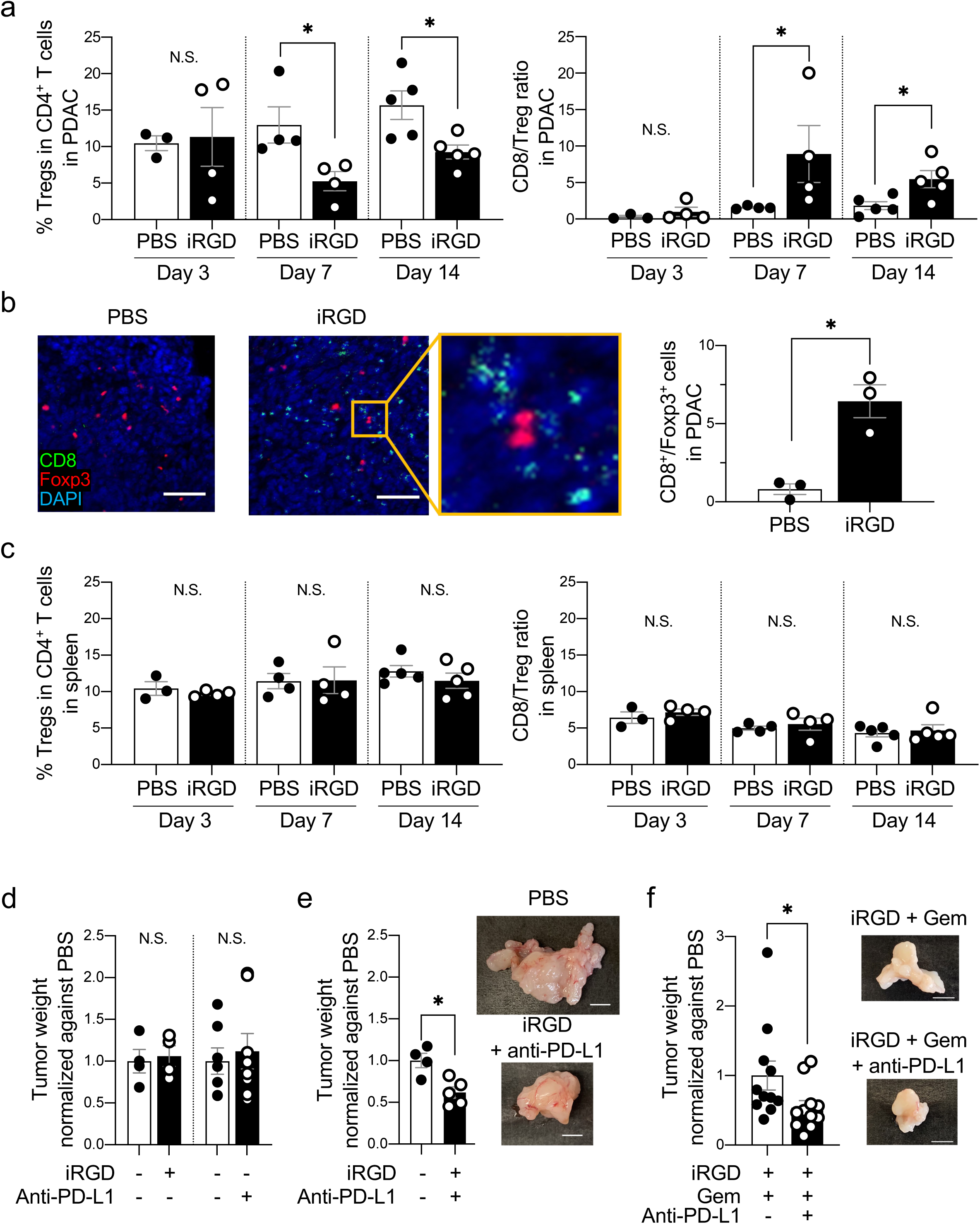
iRGD depletes Tregs and expands CD8^+^ T cells in PDAC tissue, and enhances the efficacy of an anti-PD-L1 blockade. **a-c** C57B6129SF1/J hybrid mice bearing KPC-derived orthotopic PDAC tumors were intravenously treated with PBS or iRGD 3 times a week for 2 weeks. Tumors and spleens were collected 3, 7 and 14 days after the treatment was started. The proportion of CD4^+^ CD25^+^ Tregs among CD4^+^ T cells in the tumor (**a**, left panel) or the spleen (**c**, left panel), and the ratio of CD8^+^ T cells over Tregs (CD8/Treg ratio) in the tumor (**a**, right panel) or the spleen (**c**, right panel) were analyzed by flow cytometry. n = 3 (day 3, PBS), 4 (day3, iRGD; day 7) or 5 (day 14) per group. Representative images of IF performed on the PDAC tissue harvested after 14 days of treatment are shown in (**b**). Green, CD8^+^ T cells; red, Foxp3^+^ Tregs; blue, DAPI. Scale bars, 50 μm. The bar diagram on the right shows the CD8/Treg ratio, which was quantified by counting the cells in 5 random high-power fields per section for 3 mice per group. Statistical analysis, Welch’s *t* test (for n = 3) or Mann-Whitney U test (for n ≧ 4); *p* = 0.8454 (**a**, left, day 3), *p* = 0.0286 (**a**, left, day 7), *p* = 0.0317 (**a**, left, day 14), *p* = 0.3656 (**a**, right, day 3), *p* = 0.0286 (**a**, right, day 7), *p* = 0.0317 (**a**, right, day 14), *p* = 0.0243 (**b**), *p* = 0.5195 (**c**, left, day 3), *p* = 0.8857 (**c**, left, day 7), *p* = 0.5476 (**c**, left, day 14), *p* = 0.4696 (**c**, right, day 3), *p* = 0.3429 (**c**, right, day 7), *p* = 0.6905 (**c**, right, day 14). **d-f** KPC-derived orthotopic PDAC mice were treated with PBS, iRGD, anti-PD-L1 Ab, or iRGD + anti-PD-L1 mAb (**d, e**) or iRGD + anti-PD-L1 mAb with or without Gem (**f**) 3 times a week for 2 weeks. The weight of the tumors at the end of the study are shown. Representative tumor images are shown to the right. Scale bars, 5 mm. n = 4, 7 or 8 per arm (**d**); n = 4 or 5 per arm (**e**); n = 11 per arm (**f**). Statistical analysis, Mann-Whitney U test; *p* > 0.9999 (**d**, left), *p* = 0.7789 (**d**, right), *p* = 0.0317 (**e**), *p* = 0.0192 (**f**). Error bars, mean ± standard error; \**p* < 0.05; N.S., not significant.

### iRGD in combination with an ICB reduces tumor growth

The ability of iRGD to reduce PDAC-resident Tregs prompted us to test whether iRGD would improve the efficacy of ICBs against PDAC. We elected to use a monoclonal Ab (mAb) against programmed cell death ligand-1 (PD-L1) given the high expression of PD-L1 in our orthotopic PDAC model (Supplementary Fig. 9). While iRGD or the anti-PD-L1 mAb alone did not affect tumor growth (Fig. 1d), iRGD in combination with the anti-PD-L1 mAb led to a significant decrease in tumor volume (Fig. 1e). Adding the anti-PD-L1 mAb to the iRGD + Gem regimen also enhanced the anti-tumor effect (Fig. 1f).

### PDAC-resident Tregs express the αvβ5 integrin

The expression of αv integrins along with NRP-1 is essential for iRGD to target cells effectively^21, 25^. IF showed that αvβ5 integrin^+^ and NRP-1^+^ Tregs were indeed present in KPC-derived orthotopic PDAC tumors (Fig. 2a). Flow cytometry showed that αvβ5 integrin was detectably expressed on approximately 20% of the Tregs (Fig. 2b and 2c). Interestingly, this integrin was rarely expressed on splenic Tregs (Supplementary Fig. 10) suggesting that selective expression of αvβ5 on PDAC-resident Tregs allowed iRGD to induce tumor-specific Treg reduction. NRP-1 was expressed on both PDAC and splenic Tregs consistent with NRP-1 being a general mouse Treg marker^32^. The expression of αvβ5 integrin and NRP-1 was significantly lower on other CD4^+^ T cells and almost undetectable on CD8^+^ T cells. Of note, αvβ5 integrin^+^ Tregs were also detected in the tumors of human PDAC patients, but minimally in their spleens (Fig. 2d and 2e). There were also slightly more NRP-1^+^ Tregs in the tumors than in the spleens, but the difference did not reach significance with the limited number of samples analyzed.

**Figure 2.**
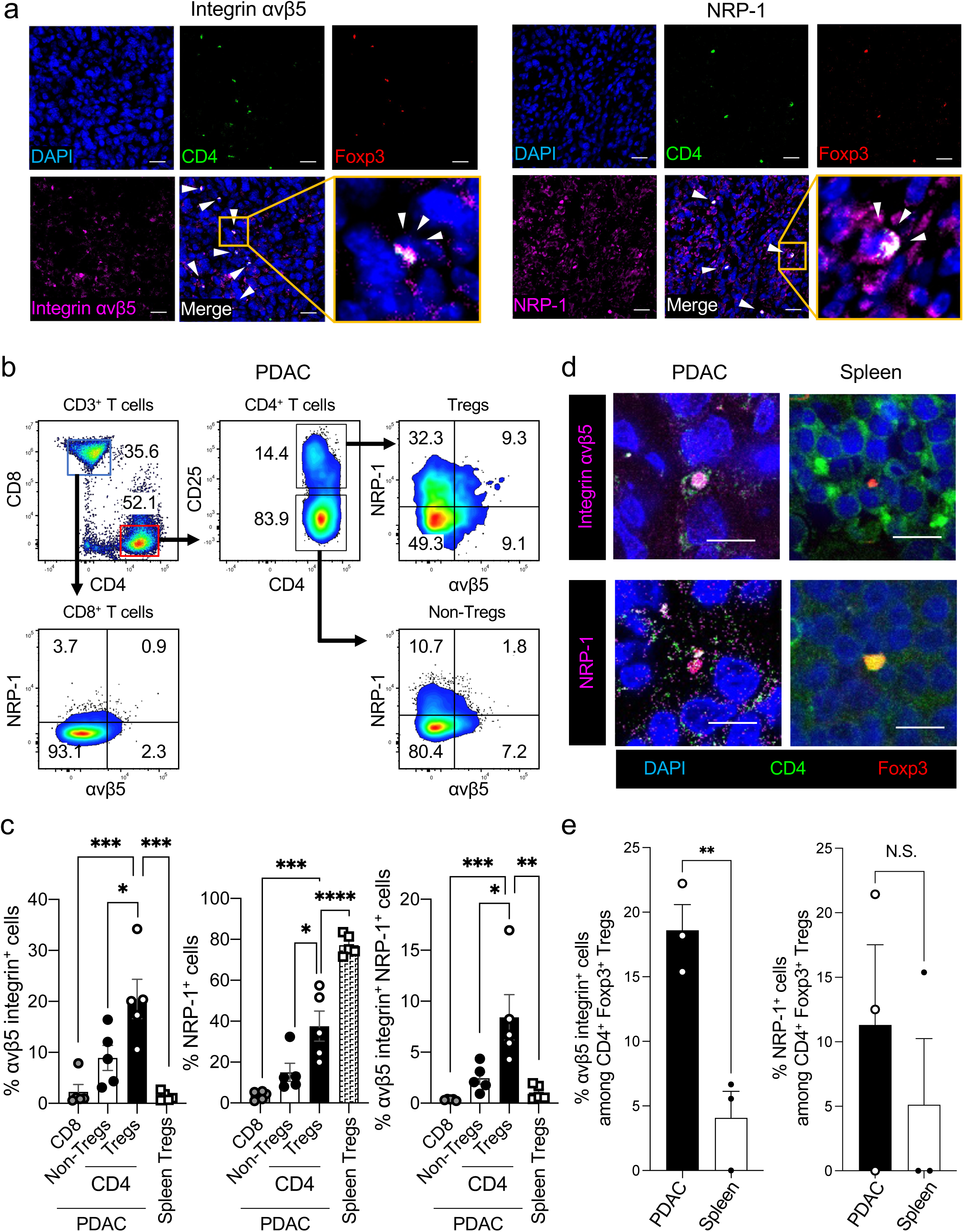
PDAC-resident Tregs express the αvβ5 integrin. **a** Representative confocal images of αvβ5 integrin^+^ CD4^+^ Foxp3^+^ T cells (left) and NRP-1^+^ CD4^+^ Foxp3^+^ T cells (right) in the PDAC tissue of KPC-derived orthotopic PDAC mice. Arrows indicate αvβ5 integrin^+^ or NRP-1^+^ Tregs (magenta). Green, CD4; red, Foxp3; blue, DAPI. The boxed areas are magnified. Scale bars, 20 μm. **b** A representative flow cytometry analysis showing the proportion of CD8^+^ T cells, CD4^+^ CD25^neg^ T cells (non-Tregs), CD4^+^ CD25^+^ Tregs that are positive for αvβ5 integrin, NRP-1, or both in KPC-derived orthotopic PDAC tumors. **c** Bar diagrams that summarize the findings from (**b**) and Supplementary Fig. 10. n = 5 per group. **d** Representative confocal images of αvβ5 integrin^+^ Foxp3^+^ CD4^+^ Tregs and NRP-1^+^ CD4^+^ Tregs in human PDAC and spleen. Magenta, αvβ5 integrin or NRP-1; green, CD4; red, Foxp3; blue, DAPI. Scale bars, 20 μm. **e** The number of αvβ5 integrin-positive and NRP-1-positive CD4^+^ Foxp3^+^ Tregs was counted under a confocal microscope and the % positivity was calculated. n = 3. Statistical analysis, one-way ANOVA (**c**) and Welch’s test (**e**); *p* = 0.0003 (**c,** left, CD8 vs Tregs), *p* = 0.0169 (**c,** left, Non-Tregs vs Tregs), *p* = 0.0002 (**c,** left, Tregs vs Spleen Tregs), *p* = 0.0003 (**c,** center, CD8 vs Tregs), *p* = 0.0123 (**c,** center, Non-Tregs vs Tregs), *p* < 0.0001 (**c,** center, Tregs vs Spleen Tregs), *p* = 0.0008 (**c,** right, CD8 vs Tregs), *p* = 0.0109 (**c,** right, Non-Tregs vs Tregs), *p* = 0.0019 (**c,** right, Tregs vs Spleen Tregs), *p* = 0.0071 (**e,** left), *p* = 0.4858 (**e,** right). Error bars, mean ± standard error. \**p* < 0.05; \*\**p* < 0.01; \*\*\**p* < 0.001; \*\*\*\**p* < 0.0001; N.S., not significant.

### PDAC cells facilitate the induction of αvβ5 integrin^+^ Tregs

The results above suggest that the αvβ5 integrin is a tumor-specific marker for PDAC-resident Tregs, and that the tumor microenvironment is critical for the induction of αvβ5 integrin^+^ Tregs. To test this possibility, we magnetically isolated CD4^+^ T cells from mouse spleens and cultured them on a monolayer of KPC-derived PDAC cells for 3 days. The presence of PDAC cells greatly enhanced the expression of αvβ5 integrin^+^ CD4^+^ CD25^+^ Tregs (Fig. 3a, top panel). The Tregs consistently expressed NRP-1 regardless of the presence of PDAC cells (Fig. 3a, bottom panel). Non-Treg CD4^+^ CD25^neg^ T cells expressed significantly less αvβ5 integrin (and NRP-1) than Tregs in the presence and absence of PDAC cells (Fig. 3b).

**Figure 3.**
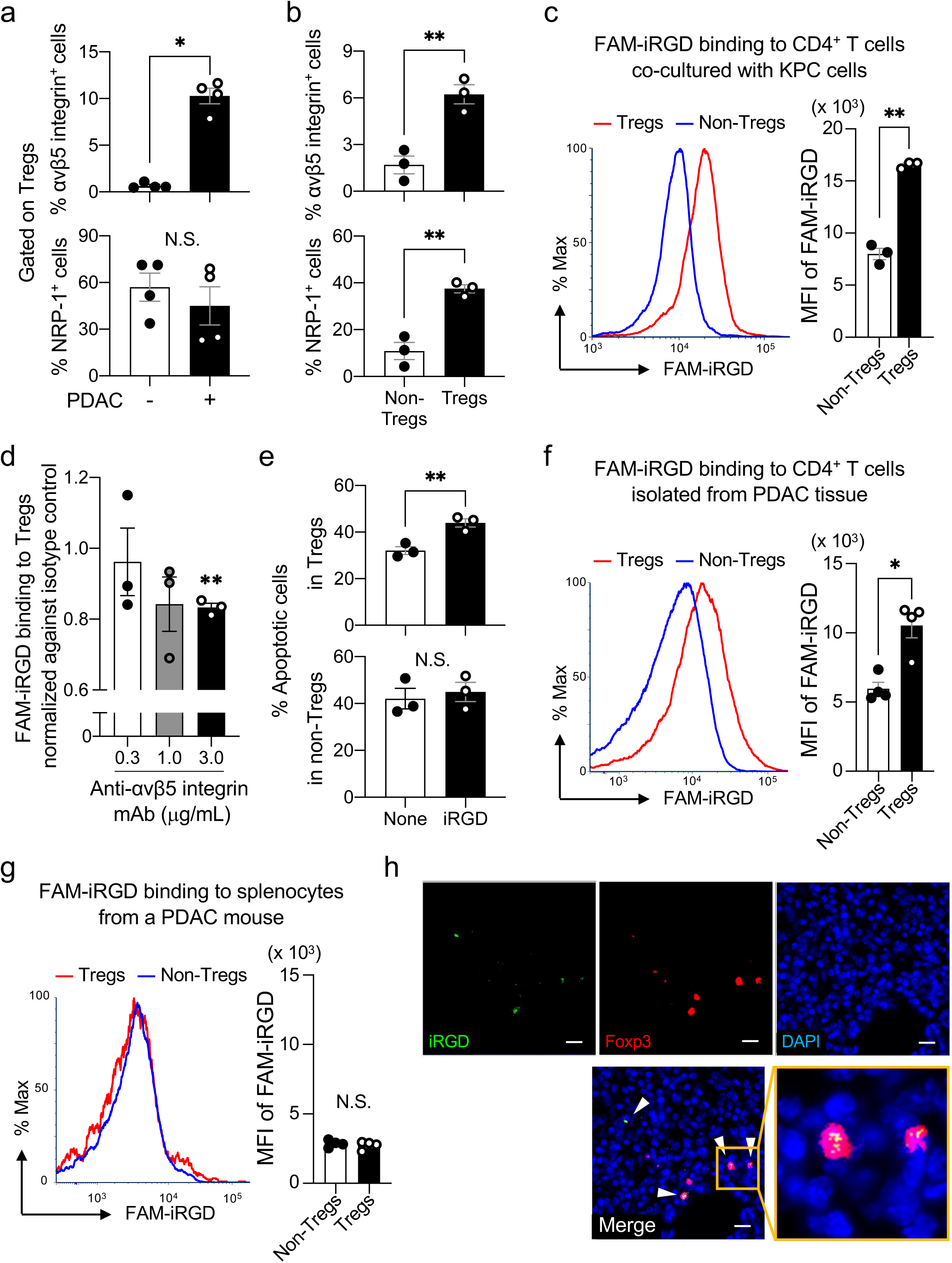
iRGD binds to αvβ5 integrin^+^ Tregs induced in the presence of PDAC cells. **a-e** CD4^+^ T cells isolated from the spleens of healthy C57B6129SF1/J hybrid mice were expanded for 3 days *in vitro* in the presence of KPC-derived PDAC cells. Flow cytometry was performed for subsequent analyses. (**a**) Expression of αvβ5 integrin and NRP-1 on CD4^+^ CD25^+^ Tregs expanded with or without the PDAC cells. n = 4 per group. (**b**) Expression of αvβ5 integrin and NRP-1 on CD4^+^ CD25^+^ Tregs and non-Treg CD4^+^ CD25^neg^ T cells expanded with the PDAC cells. n = 3 per group. (**c**) FAM-iRGD binding to the non-Tregs (blue line) and Tregs (red line) shown in (**b**). The bar diagram summarizes the median fluorescence intensity (MFI) from 4 independent experiments. (**d**) Dose-dependent inhibition of FAM-iRGD binding by an anti-αvβ5 integrin blocking Ab to Tregs that were expanded in the presence of PDAC cells. Values were normalized against isotype control. n = 3 per group. Statistical analysis was performed between the isotype control and anti-αvβ5 integrin values. (**e**) Tregs and non-Tregs were expanded in the presence of PDAC cells with or without iRGD. Apoptosis was quantified by measuring annexin V and 7-AAD double positive cells by flow cytometry. n = 3. **f, g** *In vitro* binding of FAM-iRGD to non-Treg CD4^+^ CD25^neg^ T cells (blue line) and CD4^+^ CD25^+^ Tregs (red line) isolated from the PDAC tissue (**f**) or the spleen (**g**) of KPC-derived orthotopic PDAC mice. The bar diagrams summarize the MFI from 4 independent experiments. **h** Representative confocal images of Foxp3^+^ T cells (red) in the PDAC tissue of KPC-derived orthotopic PDAC mice that received an intravenous injection of FAM-iRGD (green). Blue, DAPI. Arrows indicate Tregs positive for iRGD. The boxed area is magnified. Scale bars, 20 μm. Statistical analyses; Mann-Whitney U test (**a**, **c**, **f**, **g**), Welch’s *t* test (**b**, **e**), and one sample Wilcoxon signed rank test (**d**); *p* = 0.0286 (**a**, top), *p* = 0.3429 (**a**, bottom), *p* = 0.0058 (**b**, top), *p* = 0.0086 (**b**, bottom), *p* = 0.0017 (**c**), *p* = 0.7272 (**d**, 0.3), *p* = 0.1766 (**d**, 1.0), *p* = 0.0052 (**d**, 3.0), *p* = 0.0087 (**e**, top), *p* = 0.6583 (**e**, bottom), *p* = 0.0286 (**f**), *p* = 0.6857 (**g**). Error bars, mean ± standard error; \**p* < 0.05; \*\**p* < 0.01; N.S., not significant.

### iRGD targets PDAC-resident Tregs

Fluorescein (FAM)-labeled iRGD (FAM-iRGD) bound significantly more effectively to Tregs than non-Tregs that were expanded on PDAC cells *in vitro* (Fig. 3c). The FAM-iRGD binding to the Tregs was dependent on the αvβ5 integrin because a blocking Ab inhibited the binding in a dose-dependent manner (Fig. 3d). Interestingly, adding iRGD in the co-culture increased the apoptosis of Tregs without affecting non-Tregs (Fig. 3e; Supplementary Fig. 11), providing a potential mechanism of the iRGD-induced depletion of Tregs in PDAC. FAM-iRGD also bound significantly more effectively to Tregs than to non-Tregs that were isolated from KPC-derived orthotopic PDAC tumors (Fig. 3f). FAM-iRGD did not bind to Tregs or non-Tregs harvested from the spleens of KPC-derived PDAC mice (Fig. 3g). In line with these findings, FAM-iRGD injected into the tail vein of KPC-derived PDAC mice accumulated in Tregs in PDAC tissue (Fig. 3h).

### αvβ5 integrin^+^ Tregs are induced from naïve CD4^+^ T cells

Tumor-resident Tregs are believed to originate from two major sources, induced Tregs (iTregs) that differentiate in the periphery from naïve CD4^+^ T cells and natural Tregs (nTregs) that develop in the thymus^33, 34^. To understand the origin of αvβ5 integrin^+^ Tregs, we first tested whether they can be induced as iTregs from naïve CD4^+^ T cells, which were magnetically enriched from mouse splenocytes (Fig. 4, top row). The pool contained a small number of CD4^+^ Foxp3^+^ T cells, which were presumably residual nTregs. These cells had high expression of NRP-1 compared to the naïve CD4^+^ T cells (Supplementary Fig. 12a). Both the naïve CD4^+^ T cells and CD4^+^ Foxp3^+^ T cells lacked αvβ5 integrin. In line with previous reports^35^, T-cell receptor (TCR) stimulation by treating the CD4^+^ T cell pool with anti-CD3/CD28 bispecific Abs^36^ combined with TGF-β1 led to the expansion of CD4^+^ Foxp3^+^ iTregs (Fig. 4, middle row). Over 20% of the resulting iTregs expressed the αvβ5 integrin, while only a minor population of CD4^+^ Foxp3^neg^ non-Tregs expressed this integrin. The iTregs expressed more NRP-1 than the non-Tregs (Supplementary Fig. 12b). TCR stimulation alone did not lead to the development of iTregs from naïve CD4^+^ T cells as evidenced by the fact that >90% of the cells remained negative for Foxp3 (Fig. 4, bottom row, blue box). Only 3-4% of the CD4^+^ Foxp3^neg^ T cells expressed αvβ5 integrin. The results indicate that αvβ5 integrin^+^ Tregs induced from naïve CD4^+^ T cells arise as a subpopulation of iTregs. Interestingly, some of the CD4^+^ Foxp3^+^ T cells (Fig. 4, bottom row, red box) that were initially negative for αvβ5 integrin were found to express the integrin after TCR stimulation alone, suggesting that nTregs may also be a potential source for αvβ5 integrin^+^ Tregs. A significant portion of the nTregs also expressed NRP-1 (Supplementary Fig. 12c).

**Figure 4.**
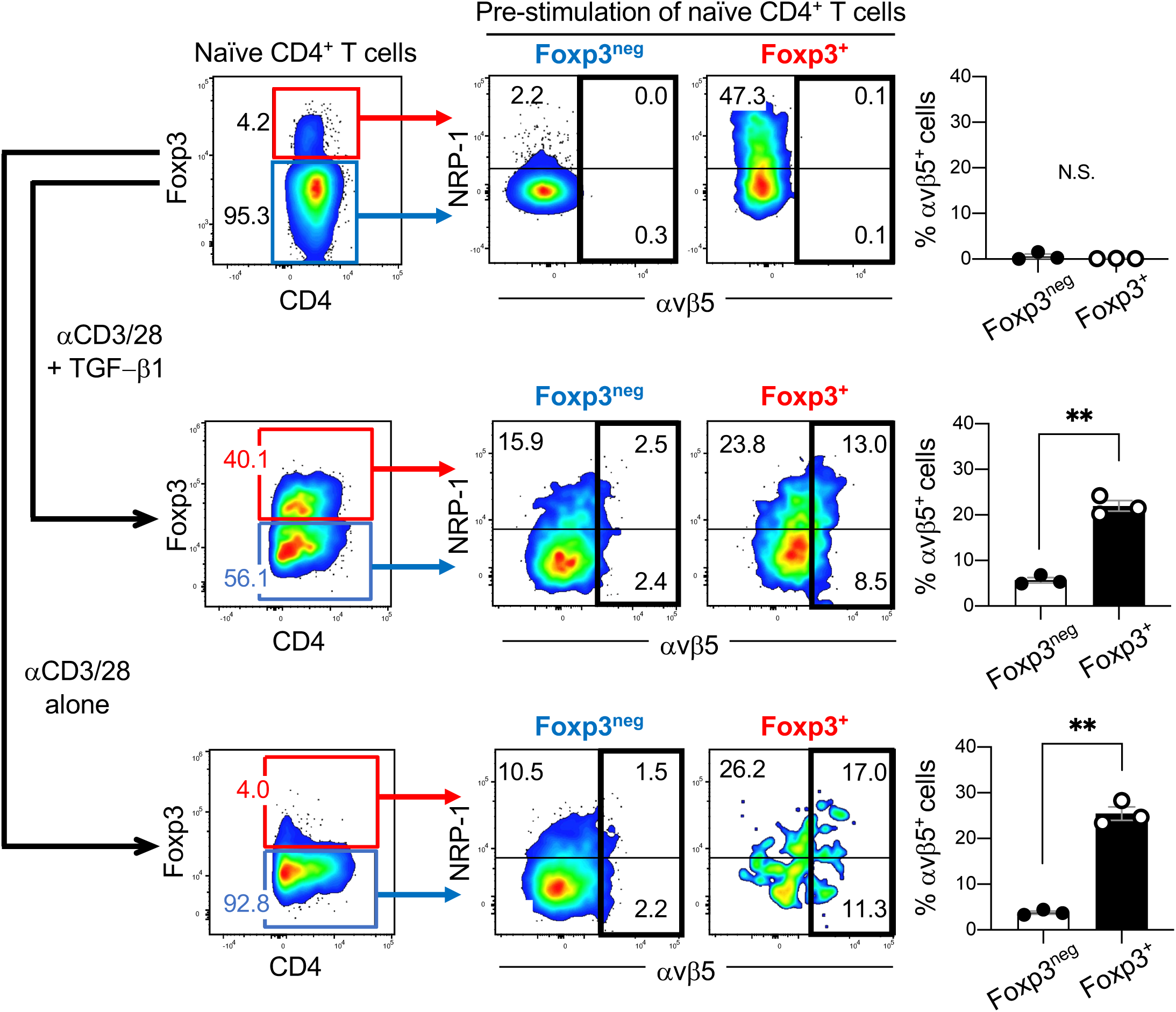
αvβ5 integrin^+^ Tregs are induced from naïve CD4^+^ T cells. Naïve CD4^+^ T cells were isolated from the spleens of healthy C57B6129SF1/J hybrid mice by magnetically removing CD4^neg^ T cells and CD25^+^ T cells. The pool enriched for naïve CD4^+^ T cells was cultured *in vitro* with anti-CD3/CD28 Abs in the presence or absence of TGF-β1 for 3 days, and analyzed for αvβ5 integrin and NRP-1 expression by flow cytometry. (**Top row)** Naïve CD4^+^ T cells (blue box) enriched from mouse splenocytes. A minor population of CD4^+^ Foxp3^+^ T cells was present (red box). (**Middle row**) Treating the pool in the top row with anti-CD3/CD28 Abs and TGF-β1 yielded approximately 40% of CD4^+^ Foxp3^+^ T cells (red box) and 56% of CD4^+^ Foxp3^neg^ T cells (blue box). (**Bottom row**) Treating the pool in the top row with anti-CD3/CD28 Abs alone did not change the proportion of the CD4^+^ T cells. Nearly 95% of the cells remained negative for Foxp3 (blue box). Representative dot plots showing the proportion of CD4^+^ Foxp3^+^ T cells (left panels) and the expression of αvβ5 integrin and NRP-1 on the indicated population are presented. The bar diagrams summarize the proportion of αvβ5^+^ cells in the indicated population. n = 3 per study. Statistical analysis, Welch’s *t* test; *p* = 0.4115 (top), *p* = 0.0013 (middle), *p* = 0.0032 (bottom). Error bars, mean ± standard error; \*\**p* < 0.01; N.S., not significant.

### Induction of αvβ5 integrin expression on iTregs is not dependent on TGF-β1

Based on the above data, it seemed possible that TGF-β1 was an inducer of αvβ5 integrin^+^ iTregs. TGF-β1 is a well-accepted inducer of integrin expression on various cells^37–40^ and we recently found that TGF-β1 secreted by CAFs (and PDAC cells) induced αvβ5 integrin expression on PDAC cells^25^. As expected, LY2157299, an inhibitor of TGF-β receptor type 1 (TGF-βR1)^25^, suppressed the development of iTregs from naïve CD4^+^ T cells in a dose-dependent manner (Supplementary Fig. 13a). However, the inhibitor did not affect the proportion of the αvβ5-positive (or NRP-1-positive) iTregs (Supplementary Fig. 13b). Thus, TGF-β1 is indeed a critical inducer of iTregs, but does not specifically induce the αvβ5-positive population among the iTregs.

### αvβ5 integrin is expressed in response to TCR stimulation

Further analysis revealed that αvβ5 integrin expression correlated with CD25, which is expressed on CD4^+^ T cells upon TCR stimulation^41^. Nearly 30% of the CD4^+^ CD25^+^ Foxp3^+^ iTregs that were induced from naïve CD4^+^ T cells by TCR stimulation and TGF-β1 expressed the αvβ5 integrin, while almost none of the CD4^+^ CD25^neg^ Foxp3^+^ T cells expressed the integrin (Fig. 5a, top row). Although the proportion was significantly less than the iTregs, some of the CD4^+^ CD25^+^ Foxp3^neg^ T cells derived from naïve CD4^+^ T cells also expressed the αvβ5 integrin (Fig. 5a, bottom row). CD4^+^ CD25^neg^ Foxp3^neg^ T cells remained negative for αvβ5 integrin expression. Similar to the expression profile of the αvβ5 integrin, NRP-1 expression was significantly higher on CD25^+^ cells (Supplementary Fig. 14a and 14b). CD4^+^ CD25^+^ Foxp3^neg^ T cells that developed from naïve CD4^+^ T cells with TCR stimulation alone also showed minor expression of the αvβ5 integrin (Fig. 5b). The cells also expressed NRP-1 to some extent (Supplementary Fig. 14c). Moreover, αvβ5 integrin^+^ CD4^+^ CD25^+^ Foxp3^+^ iTregs were dose-dependently induced by an anti-CD3 Ab (Fig. 5c), which provides TCR-mediated T cell activation signals^42^. These results suggest that TCR stimulation is likely a key inducer of αvβ5 integrin (and NRP-1) expression on iTregs.

**Figure 5.**
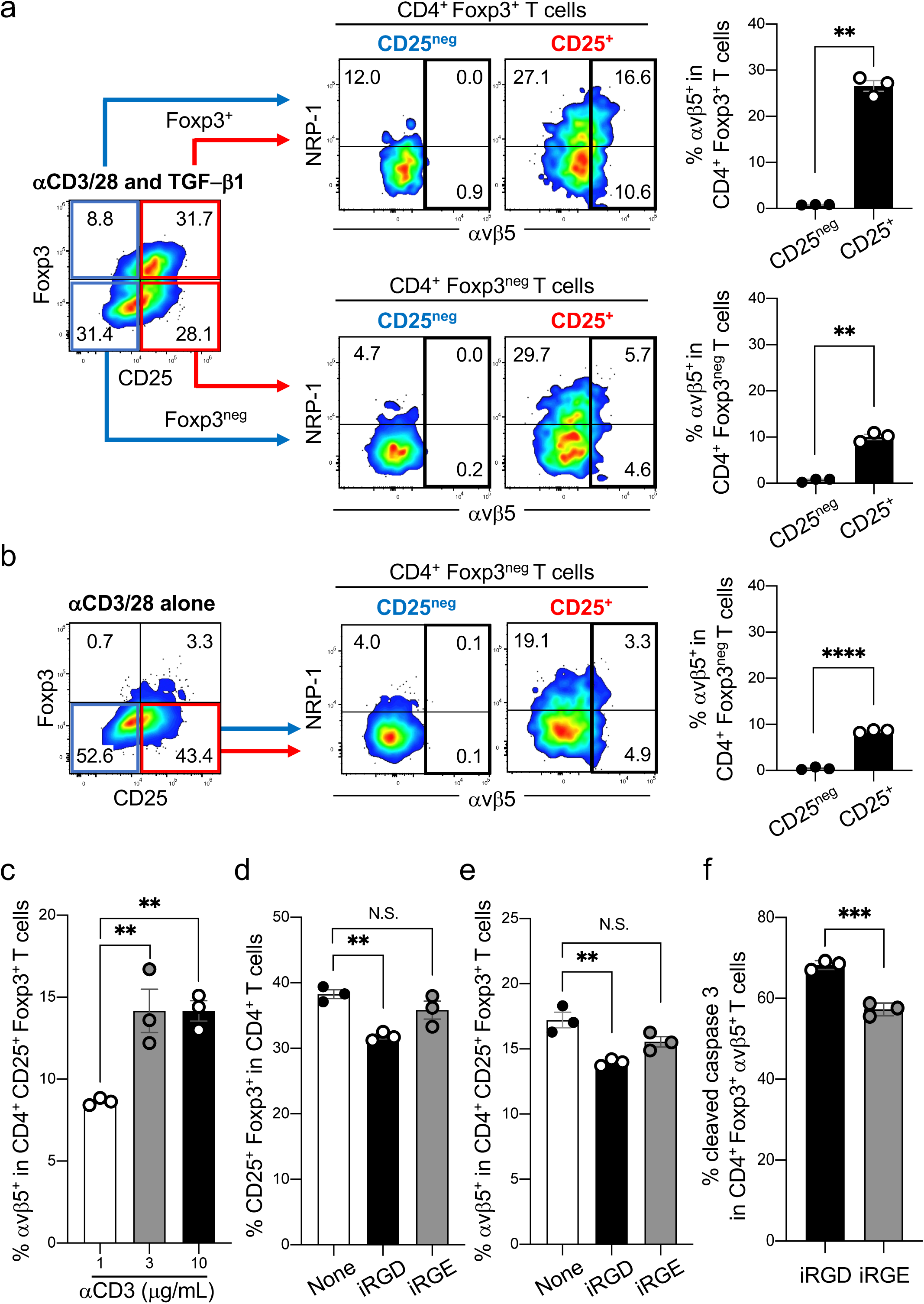
αvβ5 integrin expression is induced in response to TCR stimulation and correlates with CD25 expression. **a, b** Naïve CD4^+^ T cells isolated from healthy mouse spleens were expanded in the presence of anti-CD3/CD28 Abs and TGF-β1 (**a**) or anti-CD3/CD28 Abs alone (**b**) for 3 days. The resulting populations were gated based on Foxp3 and CD25 expression (left panels). αvβ5 integrin and NRP-1 expression on Foxp3^+^ cells (**a**, top row) and Foxp3^neg^ cells (**a**, bottom row; **b**) was analyzed by flow cytometry. The red and blue boxes gate CD25^+^ and CD25^neg^ cells, respectively. The bar diagrams summarize the proportion of αvβ5 integrin-positive cells in the indicated T cell population. n = 3. Statistical analysis, Welch’s *t* test; *p* = 0.0021 (**a**, top), *p* = 0.0014 (**a**, bottom), *p* < 0.0001 (**b**). **c** Flow cytometric analysis showing the proportion of αvβ5 integrin^+^ cells among CD4^+^ CD25^+^ Foxp3^+^ iTregs induced by increasing concentrations of anti-CD3 Ab. n = 3. Statistical analysis, one-way ANOVA; *p* = 0.0088 (1 vs 3), *p* = 0.0088 (1 vs 10). **d-f** Naïve CD4^+^ T cells were stimulated with anti-CD3/CD28 Abs and TGF-β1 in the absence or presence of iRGD or iRGE. Flow cytometry was performed to quantify the proportion of CD4^+^ CD25^+^ Foxp3^+^ iTregs (**d**) and αvβ5 integrin^+^ cells among the iTregs (**e**). Apoptosis of αvβ5 integrin^+^ iTregs was quantified by measuring cleaved caspase 3 using flow cytometry (**f**). n = 3. Statistical analysis, one-way ANOVA (**d**, **e**) or Welch’s *t* test (**f**); *p* = 0.0042 (**d**, None vs iRGD), *p* = 0.1774 (**d**, None vs iRGE), *p* = 0.0027 (**e**, None vs iRGD), *p* = 0.0529 (**e**, None vs iRGE), *p* = 0.001 (**f**). Error bars, mean ± standard error; \**p* < 0.05; \*\**p* < 0.01; \*\*\*\**p* < 0.0001; N.S., not significant.

iRGD reduced CD4^+^ CD25^+^ Foxp3^+^ iTregs and αvβ5-positive cells among them that were induced from naïve CD4^+^ T cells *in vitro* (Fig. 5d and 5e, Supplementary Fig. 15a and 15b). iRGD did not alter the proportion of NRP-1-positive iTregs (Supplementary Fig. 15c). The effect was dependent on the integrin-binding RGD motif because iRGE (CRGEKGPDC), an iRGD variant that lacks the RGD motif but retains the NRP-1 binding motif^21^, was ineffective. The reduction was in part due to increased apoptosis of αvβ5^+^ Tregs as indicated by elevated levels of cleaved caspase 3 (Fig. 5f; Supplementary Fig. 15d). Neither iRGD nor iRGE increased the apoptosis of CD4^+^ Foxp3^neg^ T cells (Supplementary Fig. 15e).

### nTregs express the αvβ5 integrin upon TCR stimulation

We then explored whether TCR stimulation also induces αvβ5 integrin expression on nTregs. It seemed likely given that the minor CD4^+^ Foxp3^+^ T cell population in the pool enriched for naïve CD4^+^ T cells turned positive for αvβ5 integrin upon TCR stimulation (refer to Fig. 4, bottom row). Accordingly, we prepared a pool of CD4^+^ T cells enriched for nTregs using CD4 and CD25 as selection markers. Approximately 35% was CD4^+^ CD25^+^ Foxp3^+^ nTregs in the resulting pool (Fig. 6a), which was nearly 10 times more than that in the pool enriched for naïve CD4^+^ T cells (refer to Fig. 4, top left panel). Both the nTregs and the remaining naïve CD4^+^ CD25^neg^ Foxp3^neg^ T cells were negative for the αvβ5 integrin. The nTregs expressed NRP-1 as expected (Supplementary Fig. 16a). Treating the pool with anti-CD3/CD28 Abs led to the induction of αvβ5^+^ CD4^+^ CD25^+^ Foxp3^+^ T cells (Fig. 6b, top row). These cells were most likely nTregs, and not iTregs that were induced from naïve CD4^+^ T cells, because anti-CD3/CD28 Abs alone did not induce iTregs under the same condition (refer to Fig. 4, bottom row). The expression of αvβ5 integrin was noted exclusively on nTregs, and not on CD4^+^ CD25^neg^ Foxp3^+^ T cells, which act as peripheral reservoirs of differentiated nTregs that become CD25-positive upon expansion or activation^43^. Consistent with our earlier data, CD4^+^ CD25^+^ Foxp3^neg^ T cells, which likely expanded from naïve CD4^+^ T cells upon TCR stimulation, expressed low levels of αvβ5 integrin (Fig. 6b, bottom row). NRP-1 expression was also mainly restricted to CD25^+^ cells (Supplementary Fig. 16b and 16c). These results suggest that αvβ5 integrin expression can be induced on nTregs through TCR stimulation. Of note, while the mode of expression of αvβ5 integrin and CD25 correlates to some extent, it does not completely match because the CD25^+^ nTregs prior to TCR stimulation did not express the αvβ5 integrin. Thus, taken together with the iTreg data, αvβ5 integrin may serve as an activation marker for Tregs that received TCR stimulation in the periphery.

**Figure 6.**
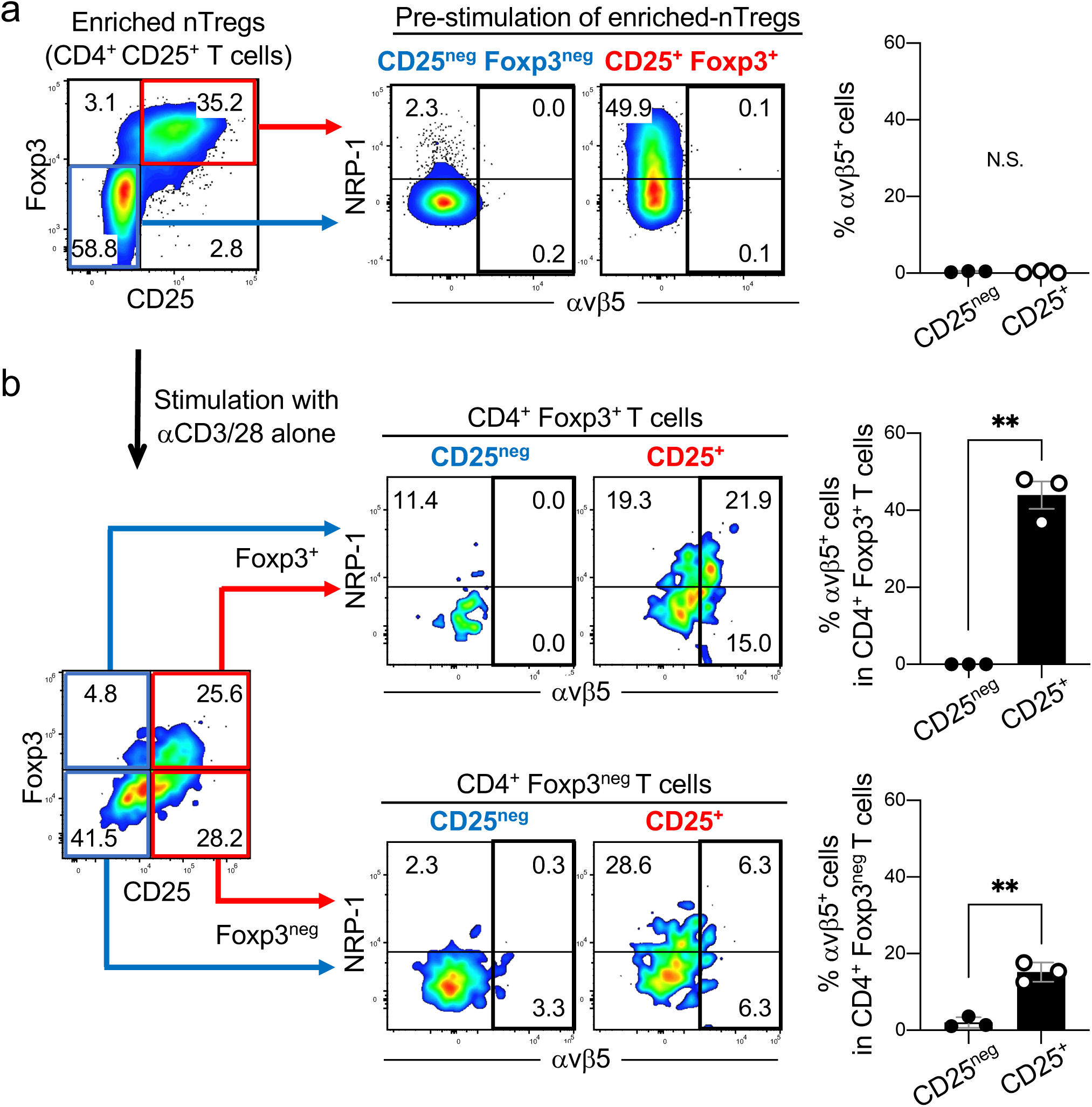
nTregs express the αvβ5 integrin in response to TCR stimulation. **a** CD4^+^ CD25^+^ Foxp3^+^ T cells (nTregs) were enriched from the spleens of healthy C57B6129SF1/J hybrid mice by magnetically removing CD4^neg^ T cells and CD25^neg^ T cells. The middle two panels show the expression of αvβ5 integrin and NRP-1 on the nTregs (red box) and naïve CD4^+^ CD25^neg^ Foxp3^neg^ T cells (blue box) analyzed by flow cytometry. The bar diagram summarizes the proportion of αvβ5 integrin^+^ cells among the two populations. n = 3. Statistical analysis, Welch’s *t* test; *p* = 0.4721. **b** The pool in (**a**) was treated with anti-CD3/CD28 Abs alone for 3 days (left panel). αvβ5 integrin and NRP-1 expression on Foxp3^+^ cells (top row) and Foxp3^neg^ cells (bottom row) was analyzed by flow cytometry. The red and blue boxes gate CD25^+^ and CD25^neg^ cells, respectively. The bar diagrams summarize the proportion of αvβ5 integrin^+^ cells among the indicated T cell populations. n = 3. Statistical analysis, Welch’s *t* test; *p* = 0.0064 (top), *p* = 0.0035 (bottom). Error bars, mean ± standard error; \*\**p* < 0.01; N.S., not significant.

### αvβ5 integrin defines a subpopulation of CCR8^+^ Tregs with enhanced immunosuppressive properties

CCR8 is one of the few markers of highly immunosuppressive tumor-resident Tregs^17, 19, 20^. Targeting CCR8 leads to tumor-specific depletion of Tregs and enhanced tumor-specific immunity, which resembles the outcome of iRGD therapy in PDAC mice. We therefore tested whether αvβ5 integrin and CCR8 have similar expression profiles. Treating naïve CD4^+^ T cells with anti-CD3/CD28 Abs and TGF-β1 led to the induction of CCR8^+^ CD4^+^ CD25^+^ Foxp3^+^ iTregs (Fig. 7a and 7b). αvβ5 integrin was expressed on approximately 25% of the CCR8^+^ iTregs and nearly all the αvβ5 integrin^+^ iTregs expressed CCR8, indicating that αvβ5 integrin expression was restricted to a subpopulation of CCR8^+^ iTregs (Fig. 7c). αvβ5 integrin was minimally expressed on CD4^+^ CD25^+^ Foxp3^neg^ non-Tregs, while CCR8 was expressed on approximately 30% of the cells, suggesting that αvβ5 integrin was more Treg-specific than CCR8. CD4^+^ CD25^neg^ Foxp3^neg^ T cells remained negative for both CCR8 and αvβ5 integrin.

**Figure 7.**
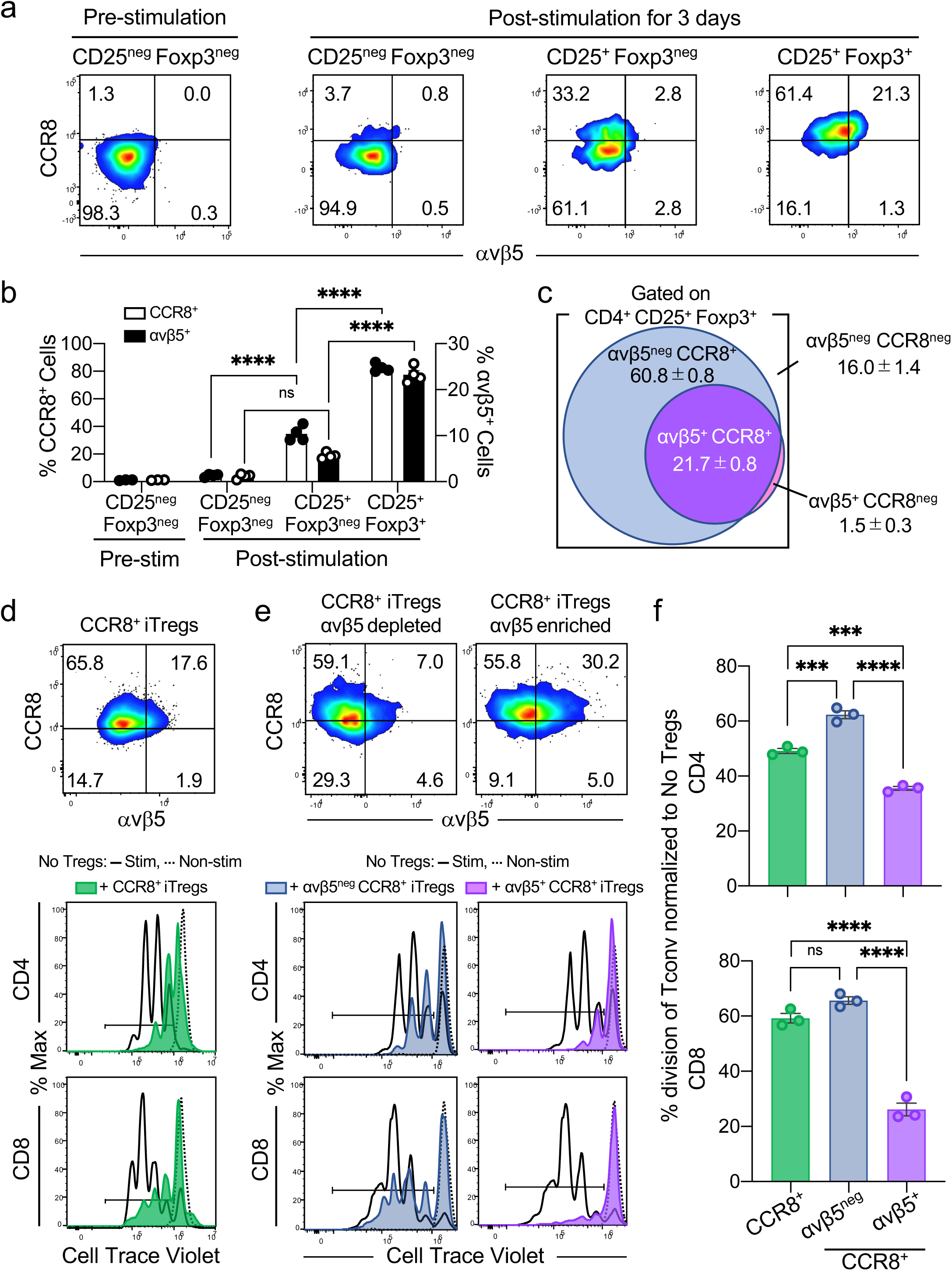
The αvβ5 integrin marks a highly immunosuppressive subpopulation of CCR8^+^ Tregs. Naïve CD4^+^ CD25^neg^ Foxp3^neg^ T cells were magnetically isolated from the spleens of healthy C57B6129SF1/J mice. The cells were stimulated with anti-CD3/CD28 Abs and TGF-β1 for 3 days to induce CD4^+^ CD25^+^ Foxp3^+^ iTregs. **a-c** Expression of CCR8 and αvβ5 integrin on the T cells before and after the stimulation. Representative dot plots from 3 or 4 separate studies are shown in (**a**). The bar diagram in (**b**) summarizes the proportion of CCR8^+^ cells (white bars) and αvβ5 integrin^+^ cells (black bars) among each T cell population in (**a**). The Venn diagram in (**c**) summarizes the proportion of iTregs that expressed CCR8 and/or αvβ5 integrin. **d-f** Treg suppression assays were performed by co-culturing iTregs and Tconv (CD4^+^ and CD8^+^) at a 1 : 4 ratio in the presence of anti-CD3/CD28 Abs (TCR stimulation) for 3 days. We used iTregs that were enriched for CCR8^+^ iTregs (**d**) or CCR8^+^ iTregs that were either depleted or enriched for αvβ5 integrin^+^ cells (**e**). The expression of CCR8 and αvβ5 integrin on the iTregs is shown in the representative dot plots. Proliferation of Tconv was analyzed by flow cytometry using Cell Trace Violet as shown in the representative histograms: Shaded, iTregs + Tconv (with TCR stimulation); black solid line, Tconv alone (with TCR stimulation); black dotted line, Tconv alone (no TCR stimulation). The bar diagrams in (**f**) summarize the values from (**d**) and (**e**) normalized to stimulated Tconv alone. n = 3. Statistical analysis, one-way ANOVA; *p* < 0.0001 (**b**, CCR8^+^, CD25^neg^ Foxp3^neg^ vs CD25^+^ Foxp3^neg^ and CD25^+^ Foxp3^neg^ vs CD25^+^ Foxp3^+^; αvβ5^+^, CD25^+^ Foxp3^neg^ vs CD25^+^ Foxp3^+^), *p* = 0.0989 (**b**, αvβ5^+^, CD25^neg^ Foxp3^neg^ vs CD25^+^ Foxp3^neg^), *p* = 0.0003 (**f**, CD4, CCR8^+^ vs αvβ5^neg^ CCR8^+^), *p* = 0.0002 (**f**, CD4, CCR8^+^ vs αvβ5^+^ CCR8^+^), *p* < 0.0001 (**f**, CD4, αvβ5^neg^ CCR8^+^ vs αvβ5^+^ CCR8^+^), *p* = 0.1048 (**f**, CD8, CCR8^+^ vs αvβ5^neg^ CCR8^+^), *p* < 0.0001 (**f**, CD8, CCR8^+^ vs αvβ5^+^ CCR8^+^ and αvβ5^neg^ CCR8^+^ vs αvβ5^+^ CCR8^+^). Error bars, mean ± standard error; \*\*\**p* < 0.001; \*\*\*\**p* < 0.0001; N.S., not significant.

Further studies revealed that the αvβ5 integrin^+^ Tregs had pronounced immunosuppressive properties. We induced iTregs from mouse naïve CD4^+^ T cells, and magnetically enriched for CCR8^+^ iTregs (Fig. 7d). Approximately 83% of the iTregs expressed CCR8, and 18% expressed both CCR8 and αvβ5 integrin. The pool effectively suppressed the proliferation of conventional CD4^+^ and CD8^+^ T cells (Tconv) in traditional immunosuppression assays. To study the contribution of αvβ5^+^ Tregs in this effect, we prepared pools of CCR8^+^ iTregs that were either magnetically enriched or depleted for αvβ5^+^ iTregs (Fig. 7e). Enriching the αvβ5^+^ population significantly enhanced the ability to suppress Tconv proliferation (Fig. 7f). The effect appeared to be more pronounced against CD8^+^ T cells than CD4^+^ T cells. In contrast, depleting αvβ5^+^ cells reduced the suppression properties. These results strongly suggest that αvβ5^+^ Tregs are highly immunosuppressive, and that they may be the functionally dominant fraction of CCR8^+^ Tregs.

## DISCUSSION

It is becoming evident that tumor-specific Treg depletion is desirable when Treg targeting is used in cancer management^44, 45^. Achieving this goal remains a challenge given the insufficient tumor specificity and cell type specificity of Treg-associated markers^46, 47^. CCR8 provides a possible opportunity to deplete functionally activated Tregs in a tumor-selective manner^16–19^. However, the functional role of CCR8 in the immunosuppressive properties of Tregs is unknown, and the efficacy and safety of CCR8-targeted therapy in human cancer patients remains unclear despite the multiple clinical trials that are underway (e.g., NCT05537740, NCT05518045, NCT04895709)^48–50^[REFs]. Our study identifies the αvβ5 integrin as a marker for highly functional tumor-resident Tregs providing a new opportunity to selectively target the Tregs for improved tumor therapy.

iRGD therapy of PDAC mice significantly increased the CD8/Treg ratio in the tumors. The expansion of CD8^+^ T cells was likely secondary to Treg depletion, rather than a direct effect on CD8^+^ T cells by iRGD, because the iRGD receptors, αvβ5 integrin and NRP-1, were not detected on the CD8^+^ T cells. These data suggest that iRGD may reverse some of the immunosuppressive features in the PDAC tumor microenvironment. Because iRGD specifically targets tumors, this effect was likely to be limited to tumors and could be useful in tumor immunotherapy. Consistent with previous studies^21, 25^, iRGD monotherapy did not inhibit tumor growth in the present PDAC study. However, the favorable change in the CD8/Treg ratio induced by iRGD treatment suggested that it might prime the tumors to stimulate tumor immunity. That the improved CD8/Treg ratio alone did not reduce tumor growth may be due to insufficient activation and/or exhaustion of the expanded CD8^+^ T cells caused by the high PD-L1 expression in the PDAC model we used. PD-1 expressed on CD8^+^ T cells inhibits naïve-to-effector differentiation of the cells and induces exhaustion after differentiation^51, 52^. Thus, blocking the PD-1/PD-L1 axis is often a necessary step to gain enhanced and prolonged effector function of CD8^+^ T cells against cancers, or infections^53–55^. Indeed, we found that iRGD and PD-L1 blockade were synergistic; the combination therapy inhibited the growth of the PDAC tumors which were refractory to both iRGD alone and anti-PD-L1. This result is in line with a previous study showing that the delivery of cytotoxic CD8^+^ T cells into PD-L1^high^ gastric tumors using iRGD only resulted in a moderate anti-tumor effect, but genetic deletion of PD-1 in the CD8^+^ T cells significantly enhanced the therapeutic efficacy^56^.

Our finding that TCR stimulation induced αvβ5 integrin expression on Tregs suggests a link between antigen recognition and αvβ5 expression, and that this may be a useful marker of antigen-specific Tregs. Induction by antigen recognition may explain why Tregs express the integrin in the presence of PDAC cells *in vitro* and *in vivo*. Antigen-specific Tregs have been assumed to be iTregs derived from naïve CD4^+^ T cells, but recent studies have revealed that nTregs can also be a source if proper stimuli are provided^57, 58^. Indeed, our data show that αvβ5 integrin^+^ Tregs can be induced from naïve CD4^+^ T cells through the iTreg route, and also from already differentiated nTregs. The fact that resting nTregs prior to TCR stimulation lacked αvβ5 integrin expression suggests that the integrin is not a pan-Treg marker but marks the activation status of Tregs. Unlike CD25 and adhesion molecules such as lymphocyte function-associated antigen-1 (also known as the αLβ2 integrin), which can be expressed on T cells upon TCR stimulation^42, 59^ αvβ5 integrin appears to be tightly linked to Tregs given its abundance on CD4^+^ CD25^+^ Foxp3^+^ iTregs compared to CD4^+^ CD25^+^ Foxp3^neg^ T cells after TCR stimulation.

Antigen-specific Tregs are highly activated, and express various chemokines, cytokines, and their receptors^59, 60^ One of them is the chemokine receptor CCR8, which is expressed on immunosuppressive Tregs that have been activated upon TCR stimulation^18–20^. CCR8^+^ Tregs are enriched in the tumor, and have become a potentially important therapeutic target to achieve tumor-specific Treg depletion^16–19^. Our finding that αvβ5 integrin^+^ Tregs are a functionally dominant fraction of CCR8^+^ Tregs suggests that αvβ5 integrin-targeted therapy may provide a more effective way to deplete functional tumor-resident Tregs. In addition, αvβ5 integrin expression appeared to be more restricted than CCR8 to Tregs that are most relevant to CD8^+^ T cell suppression, suggesting that iRGD therapy, which targets αvβ5, may provide an opportunity to further improve the tumor specificity of Treg-targeted therapy. The fact that αvβ5 integrin^+^ Tregs were highly effective in suppressing CD8^+^ T cells provides a plausible explanation for the significant expansion of CD8^+^ T cells in PDAC we achieved with iRGD treatment. Our data show that iRGD induces apoptosis of αvβ5 integrin^+^ Tregs in an RGD-dependent manner, possibly by interfering with αvβ5-dependent survival signals. Thus, it is likely that the iRGD-induced expansion of the CD8^+^ T cell population is secondary to release of the Treg-mediated suppression of CD8^+^ T cells.

The functional significance of αvβ5 integrin in Tregs remains to be elucidated. Tregs are thought to suppress the proliferation and functions of T cells through direct cell-cell contact, and indirectly by modulating antigen presenting cells^59, 61, 62^. Given the role of αvβ5 integrin as an adhesion molecule^63^, it is reasonable to speculate that the integrin enhances Treg engagement with the responder cells. αvβ5 may also provide an additional benefit by enhancing Treg survival through its known role in protecting cells from extrinsic apoptosis^64, 65^. iRGD appears to reverse these effects, which may be therapeutically useful.

In summary, we report a new way of reversing Treg-mediated suppression of tumor immunity. It is based on tumor-specific elimination of tumor Tregs achieved by targeting the αvβ5 integrin expressed on tumor-resident Tregs with the iRGD peptide. Importantly, a recent study using our humanized PDAC (huPDAC) mice suggests that the concept can be translated to human PDAC patients (Miyamura et al, submitted). The immune system of huPDAC mice is replaced by functional human immune cells, which react to antigens in a human-leukocyte antigen (HLA)-restricted manner, allowing us to study the response of human immunity against human PDAC in mice. The huPDAC tumors were found to harbor αvβ5 integrin- and NRP-1-positive human Tregs, and treatment of the mice with iRGD monotherapy increased the CD8/Treg ratio in the tumors. These findings support the argument that iRGD-mediated Treg targeting can be achieved in human PDAC patients. These findings have inspired clinical trials that are now in progress (e.g., ACTRN12623000223639).

## MATERIALS AND METHODS

### Peptides

The cyclic peptides iRGD (acetyl-CRGDKGPDC-NH_2_) and iRGE (acetyl-CRGEKGPDC-NH_2_) were synthesized in-house as previously described^21, 25^ and were also purchased from a commercial vendor (LifeTein, Hillsborough, NJ). In brief, the in-house synthesis was performed with a Liberty automatic microwave-assisted peptide synthesizer (CEM Corporation, Matthews, NC) using standard solid-phase Fmoc/t-Bu chemistry. Some of the peptides were labeled with 5(6)-carboxyfluorescein (FAM) separated with a 6-aminohexanoic acid spacer^25^.

### Tumor cells

KPC-derived PDAC cells were established from the tumors of transgenic *Kras^G12D/+^;LSL-Trp53^R172H/+^;Pdx-1-Cre* mice as previously reported^25^, and were authenticated using eighteen mouse short tandem repeat by ATCC (Manassas, VA). The cells were labeled with luciferase using a lentivirus that encoded the firefly luciferase cDNA linked to neomycin resistant cDNA via a P2A cleavage peptide (LV-Fluc-P2A-Neo; Imanis Life Sciences, Rochester, MN). The cells were cultured in Dulbecco’s modified Eagle medium with 10% fetal bovine serum (FBS) and a penicillin-streptomycin mixture. The cells tested negative for mycoplasma contamination.

### Tumor mouse models

All animal experiments were performed according to procedures approved by the Institutional Animal Care and Use Committee (IACUC) at Columbia University (New York, NY) or the University of California San Diego (UCSD, La Jolla, CA). Syngeneic PDAC mice were generated by orthotopic pancreatic injections of 5.0 x 10^5^ KPC-derived PDAC cells into 8- to 10-week-old C57B6129SF1/J hybrid mice (Jackson, Bar Harbor, ME). The mice are the offspring of a cross between C57BL/6J females and 129S1/SvImJ males. In some experiments, tumor growth was measured by luminescence imaging. The mice were anesthetized with isoflurane, and the body hair was shaved. The mice received intraperitoneal (IP) injection of 2 milligrams of luciferin (Promega Corporation, Madison, WI) and subjected to luminescence imaging using an IVIS^®^ Spectrum In Vivo Imaging System (PerkinElmer Inc, Waltham, MA). Transgenic KPC mice were maintained as described in our previous article^25^.

### Human sample collection

Human experiments were performed according to procedures approved by the Institutional Review Board at Columbia University (protocol #AAAT1231). Anonymized human specimens from scheduled operations were obtained when the pathologist determined that excess tissue was available for research purposes.

### Mouse treatment studies

Long-term treatment studies using combinations of iRGD, Gem, and Nab-P in transgenic KPC mice and subsequent immunohistochemistry (IHC) were performed as previously reported^25^. The treatment studies in orthotopic syngeneic PDAC mice were performed as follows. Eight days after orthotopic implantation of 5.0 x 10^5^ KPC-derived PDAC cells, the mice were randomized into respective treatment cohorts, such as intravenous (IV) phosphate buffered saline (PBS) alone, IV iRGD alone (300 μg/25 g), IV PBS + IP anti-mouse PD-L1 mAb (200 μg/mouse; Ultra-LEAF™ purified anti-mouse PD-L1 Ab, clone 10F.9G2; BioLegend, San Diego, CA), IV iRGD + IP anti-mouse PD-L1 mAb, IV PBS + IV Gem (4 mg/kg; MilliporeSigma, St. Louis, MO), and IV iRGD + IV GEM + IP anti-mouse PD-L1 mAb. The treatment was given 3 times a week for 2 weeks starting 8 days after tumor cell implantation. The tumors and major organs were harvested at the end of the studies on day 22, weighed, and processed for flow cytometry, IF, and IHC. The treatment studies were terminated according to the guidelines by the IACUC at Columbia University. Treatment studies in transgenic KPC mice were performed as previously described^25^.

### Isolation of immune cells from PDAC tissue

Tumors from orthotopic PDAC mice were minced into 2-4 mm pieces and added with an enzyme mix from a tumor dissociation kit (Miltenyi Biotec, Bergisch Gladbach, Germany) in Roswell Park Memorial Institute medium 1640 (Cytiva, Marlborough, MA). The tissues were dissociated using a gentle MACS Dissociator (Miltenyi Biotec) followed by a 40 min incubation at 37 °C. The samples were then centrifuged for 10 min at 3000 x *g* in the presence of debris removal solution (Miltenyi Biotec) and 10 min at 1000 x *g* in cooled PBS. The supernatant was aspirated to remove the debris. Splenocytes were prepared by gently grinding the spleen and filtrating the resulting cell suspension through a 40 µm cell strainer. Red cells were lysed with an ACK lysing buffer (Thermo Fisher Scientific, Waltham, MA). The isolated immune cells were subjected to flow cytometry for quality check and to analyze the cell populations of interest. To test FAM-iRGD binding, the isolated cells were incubated with 10 µM FAM-iRGD for 1 h at 37 °C in a binding buffer containing 1.0 % bovine serum albumin (BSA), 150 mM sodium, 1.0 mM magnesium, and 1.0 mM calcium. The binding was analyzed by flow cytometry.

### *In vitro* co-culture system of CD4^+^ T cells and KPC-derived PDAC cells

CD4^+^ T cells were magnetically isolated from the spleens of healthy C57B6129SF1/J mice using a mouse CD4^+^ T cell isolation kit (Miltenyi Biotec). The CD4^+^ T cells were cultured for 3 days with or without KPC-derived PDAC cells in the presence of low dose anti-CD3/CD28 beads (x 40 dilution; Miltenyi Biotec), TGF-β1 (5 ng/ml; Bio-Techne Corporation), and recombinant mouse interleukin-2 (mIL-2, 100 U/ml; Roche, Basel, Switzerland), and subjected to flow cytometry as described elsewhere. In some cases, 1.0 mM iRGD was added to the culture in binding buffer supplemented with mIL-2 to study the effect on apoptosis. The cells were then stained with FITC-conjugated annexin V (BioLegend) and 7-AAD (BioLegend) for 15 min at room temperature to detect apoptotic cells by flow cytometry.

### *In vivo* peptide homing assay

C57B6129SF1/J mice bearing 22 day-old orthotopic KPC tumors received an IV bolus of PBS or FAM-iRGD (300 μg/25 g). One hour later, the mice were perfused through the heart with PBS under deep anesthesia, and the tumors were collected. The tissues were stained for Foxp3 and DAPI using Abs described elsewhere, and subjected to confocal microscopy.

### Flow cytometry

Immune cells isolated from the tumor or spleen, or cells cultured *in vitro* were subjected for surface marker staining with the following reagents for 20 min at 4 °C: Brilliant Violet 650^TM^-conjugated anti-mouse CD3 Ab (clone 17A2; BioLegend), PerCP/Cyanine5.5-conjugated anti-mouse CD4 Ab (clone GK1.5; BioLegend), PE/Cyanine7-conjugated anti-mouse CD8a Ab (clone 53-6.7; BioLegend), PE-conjugated anti-mouse CD25 Ab (clone 7D4; Miltenyi Biotec), PE/Cyanine7-conjugated anti-mouse CD25 Ab (clone PC61; BioLegend), Brilliant Violet 421^TM^ or 711^TM^-conjugated anti-mouse CD304/NRP-1 Ab (clone 3E12; BioLegend), Alexa Fluor^®^ 647-conjugated anti-mouse integrin αvβ5 Ab (clone ALULA; BD Biosciences, San Jose, CA), PE-conjugated anti-mouse CD198 (CCR8) Ab (clone SA214G2; BioLegend), Zombie NIR^TM^ (BioLegend), or Aqua^TM^ Fixable Viability Kit (BioLegend). Intracellular staining of immune cells was performed using the following Abs and a Foxp3 staining buffer kit (eBioscience) according to the manufacturer’s instructions: Brilliant Violet 421^TM^-conjugated anti-Foxp3 Ab (FJK-16s; eBioscience), Pacific blue^TM^-conjugated anti-mouse Foxp3 Ab (clone MF-14; BioLegend), cleaved caspase-3 (Asp175) (5A1E) rabbit mAb (Cell Signaling, Danvers, MA), and Alexa Flour^TM^ 488-conjugated donkey anti-rabbit secondary Ab (Thermo Fisher Scientific). Tregs were defined as CD4^+^ CD25^+^ or CD4^+^ Foxp3^+^ depending on the experimental design. PD-L1 expression on KPC cells was assessed with an allophycocyanin-conjugated anti-mouse PD-L1 Ab (clone 10F.9G2; BioLegend). Flow cytometry was performed using a Cytek Aurora (Cytek, Fremont, CA) or BD LSRFortesa^TM^ (BD Biosciences), and the data were analyzed with FCS Express version 7.06.0015 (De Novo Software, Pasadena, CA) or FlowJo^TM^ v10.8 (BD Biosciences).

### Immunofluorescence

Mouse and human samples were fixed with 4% paraformaldehyde overnight, washed with PBS three times, and transferred to 30% sucrose at 4°C until the tissues sank. The tissues were then embedded in optimal cutting temperature (OCT) compound to prepare 10 μm frozen sections. The sections were stained with anti-mouse CD4 Ab (clone GK1.5; Invitrogen, Carlsbad, CA), fluorescein-conjugated anti-mouse CD8 Ab (clone YTS 105.18; Absolute Antibody, Cleveland, UK), anti-human/mouse Foxp3 Ab (clone 1054c; Novus Biologicals, Centennial, CO), anti-mouse NRP-1 Ab (R&D Systems Inc; Minneapolis, MN), Alexa Fluor^®^ 647-conjugated anti-mouse integrin αvβ5 Ab (clone ALULA; BD Biosciences), Ultra-LEAF™ purified anti-mouse PD-L1 Ab (clone 10F.9G2; BioLegend), 4’,6-diamidino-2-phenylindole (DAPI; Thermo Fisher Scientific) followed by an appropriate secondary Ab with Alexa Fluor 488, 546 or 680 (Thermo Fisher Scientific). Paraffin-embedded 4 μm sections of human tissue were obtained through the Molecular Pathology/MPSR Core at Columbia University. After treating them with IHC Antigen Retrieval Solution (00-4956-58, Invitrogen), the sections were stained with an anti-human CD4 Ab that was pre-labeled with Alexa Fluor 488 using an Ab labeling kit (Molecular Probes, A20187), anti-human/mouse Foxp3 Ab pre-labeled with Alexa Fluor 555 (Molecular Probes, A20181), Alexa Fluor® 647-conjugated anti-mouse integrin αvβ5 Ab, anti-human NRP-1 Ab (AD5-17F6; Milteny Biotec) pre-labeled with Alexa Fluor 647 (Molecular Probes, A20186), and DAPI. Images were taken with an LSM 710 confocal microscope system (Zeiss, Oberkochen, Germany). A ZEN 3.0 SR black edition software (Zeiss) was used for acquisition and analysis of the images. In some experiments, cells of interest were counted in five randomly chosen high-power fields (HPFs) per section to obtain an averaged number. The cell count was performed for three or four mice per group as indicated in each experiment.

### *In vitro* induction of αvβ5 integrin^+^ Tregs from naïve CD4^+^ T cells and nTregs

Naïve CD4^+^ T cells and nTregs were isolated from the spleens of healthy C57B6129SF1/J mice by magnetic separation using a mouse naïve CD4^+^ T cell isolation kit (Miltenyi Biotec). The T cells were stimulated with anti-CD3/CD28 beads (x 25 dilution; Miltenyi Biotec) in the presence or absence of recombinant mouse TGF-β1 (5 ng/ml; Bio-Techne Corporation, Minneapolis, MN) for 3 days. iRGD (100 µM) or iRGE (100 µM) peptide was added to the cultures in some experiments. To study the effect of TGF-β1 on the induction of αvβ5 integrin^+^ iTregs, a TGF-βR1 inhibitor (LY2157299; Selleck chemicals LLC, Houston, TX) was added at varying concentrations during the stimulation process. In some cases, naïve CD4^+^ T cells were treated with varying concentrations of plate-coated anti-mouse CD3 Ab (eBioscience) in the presence of soluble anti-mouse CD28 Ab (2 µg/ml) (eBioscience) and TGF-β1 (5 ng/ml). The cells were subjected to flow cytometry as described elsewhere.

### *In vitro* Treg suppression assay

Naïve CD4^+^ T cells isolated from the spleens of C57B6129SF1/J mice were treated with anti-CD3/CD28 beads and TGF-β1 for 3 days as described above. mIL-2 (10 ng/ml; R&D Systems Inc) was added to the culture medium from day 1 to expand iTregs. After removing dead cells with a Dead Cell Removal kit (Miltenyi Biotec), the cells were treated with a PE-conjugated anti-CD25 Ab (Miltenyi Biotec), PE-conjugated anti-CCR8 Ab (eBioscience), and/or Alexa Fluor^®^ 647-conjugated anti-β5 integrin Ab (clone ALULA; BD Biosciences) combined with an anti-PE or -AF647 magnetic separation technique (Miltenyi Biotec) to enrich for CCR8^+^ iTregs, αvβ5 integrin^+^ CCR8^+^ iTregs, or αvβ5 integrin^neg^ CCR8^+^ iTregs. Responder CD4^+^ CD25^neg^ T cells and CD8^+^ T cells (Tconv) were isolated from the spleen of healthy C57B6129SF1/J mice by magnetic separation and labeled with CellTrace^TM^ Violet (CTV; Thermo Fisher Scientific). The Tconv and iTregs were mixed at a 4 : 1 ratio and co-cultured in the presence of anti-CD3/CD28 beads (x25 dilution). Division of the Tconv was assessed by measuring the dilution of CTV by flow cytometry on day 3.

### Statistical analyses

The Mann-Whitney U test was used to compare two groups with four or more samples per group. The Welch’s test was used to compare two groups with three samples per group. One-way analysis of variance (ANOVA) was used to compare three or more groups with a normal distribution. One sample Wilcoxon signed rank test was used when the data was not assumed to be normally distributed in each group. All statistics were performed using GraphPad Prism (Ver. 8.4.3).

## ACKNOWLEDGEMENTS

We would like to thank Ms. Yoko Odagiri for assisting the experiments, and Dr. Moriya Tsuji for providing useful comments. We also thank the Flow Cytometry Microscopy and Shared Equipment Core, Molecular Pathology/MPSR Core, the Oncology Precision Therapeutics and Imaging Core, Institute for Cancer Genetics, Department of Genetics and Development, Department of Microbiology & Immunology Flow Cytometry Core and the Human Immune Monitoring Core at Columbia University. This work was supported by grants R01CA167174 (K. N. S.), R01CA155620 and the Alexandrina M. McAfee Trust Foundation (A. M. L.) and R01CA152327 (E. R.) from the National Cancer Institute of NIH, the Translational Research Grant from the Pancreatic Cancer Action Network (K. N. S.), Idea Award with Special Focus from the Department of Defense (K. N. S.), and the Research for a Cure of Pancreatic Cancer Fund (A. M. L.). K. S. was supported by the Uehara Memorial Foundation Research Fellowship (#201941078, Japan).

## AUTHOR CONTRIBUTIONS

K. N. S., A. M. L., T. H. M., K. S., Y. Kunisada, and E. R. developed the concept of the study. K. S. and Y. Kunisada performed most of the experiments. E. S. M. performed the CD8^+^ T cell staining and analysis in tissues collected from genetically engineered mice. T. H. M., N. M., and S. E. provided critical insight into the experimental design and assisted with the experiments. C. L. and Y. Kuroda assisted with flow cytometry studies. K. N. S., Y. Kunisada, K. S., and N. M. wrote the manuscript, and all the co-authors were involved in the editing process. K. N. S., A. M. L., and E. R. supervised the study.

## COMPETING INTERESTS

K. N. S. and E. R. are co-founders of Cend Therapeutics, Inc (now Lisata Therapeutics, Inc). They have ownership interest (including patents) in the company. E. R. is a member of the board of directors of the company. No potential conflicts of interest were disclosed by the other authors.

