## Supplementary Materials for "Tumor-resident regulatory T cells in pancreatic cancer express the αvβ5 integrin as a targetable activation marker"

Fig. S1. Long-term iRGD therapy increases $CD8^{+}$ T cells in the PDAC tissue in transgenic KPC mice.

Fig. S2. Long-term iRGD therapy decreases $Foxp3^{+}$ cells in the PDAC tissue in transgenic KPC mice.

Fig. S3. Rapid growth of orthotopic KPC-derived PDAC tumors in B6129SF1/J hybrid mice.

Fig. S4. iRGD monotherapy does not affect PDAC growth.

Fig. S5. Gating strategy for flow cytometric analysis of splenic and tumor-resident T cells in orthotopic KPC-derived PDAC mice.

Fig. S6. Changes in splenic Tregs and $CD8^{+}$ T cells upon iRGD monotherapy in PDAC mice.

Fig. S7. The effect of iRGD therapy on the proportion of $CD8^{+}$ T cells, Tregs, and non-Treg $CD4^{+}$ T cells in the PDAC tissue and the spleen.

Fig. S8. The proportion of Tregs and $CD8^{+}$ T cells and the CD8/Treg ratio in the spleen of normal mice and PDAC mice.

Fig. S9. KPC-derived PDAC cells express high levels of PD-L1.

Fig. S10. Representative flow cytometry analysis of αvβ5 integrin and NRP-1 expression on splenic T cells.

Fig. S11. iRGD induces apoptosis of Tregs in the presence of KPC-derived PDAC cells.

Fig. S12. NRP-1 expression on naïve $CD4^{+}$ T cells before and after TCR stimulation with or without TGF-β1.

Fig. S13. TGF-βR1 inhibitor does not affect the expression of αvβ5 integrin or NRP-1 on iTregs.

Fig. S14. Relevance of NRP-1 and CD25 expression profiles in naïve $CD4^{+}$ T cells that received TCR stimulation with or without TGF-β1.

Fig. S15. iRGD reduces $CD4^{+}$ $CD25^{+}$ $Foxp3^{+}$ iTregs in an RGD-dependent manner.

Fig. S16. NRP-1 expression on nTregs before and after TCR stimulation.

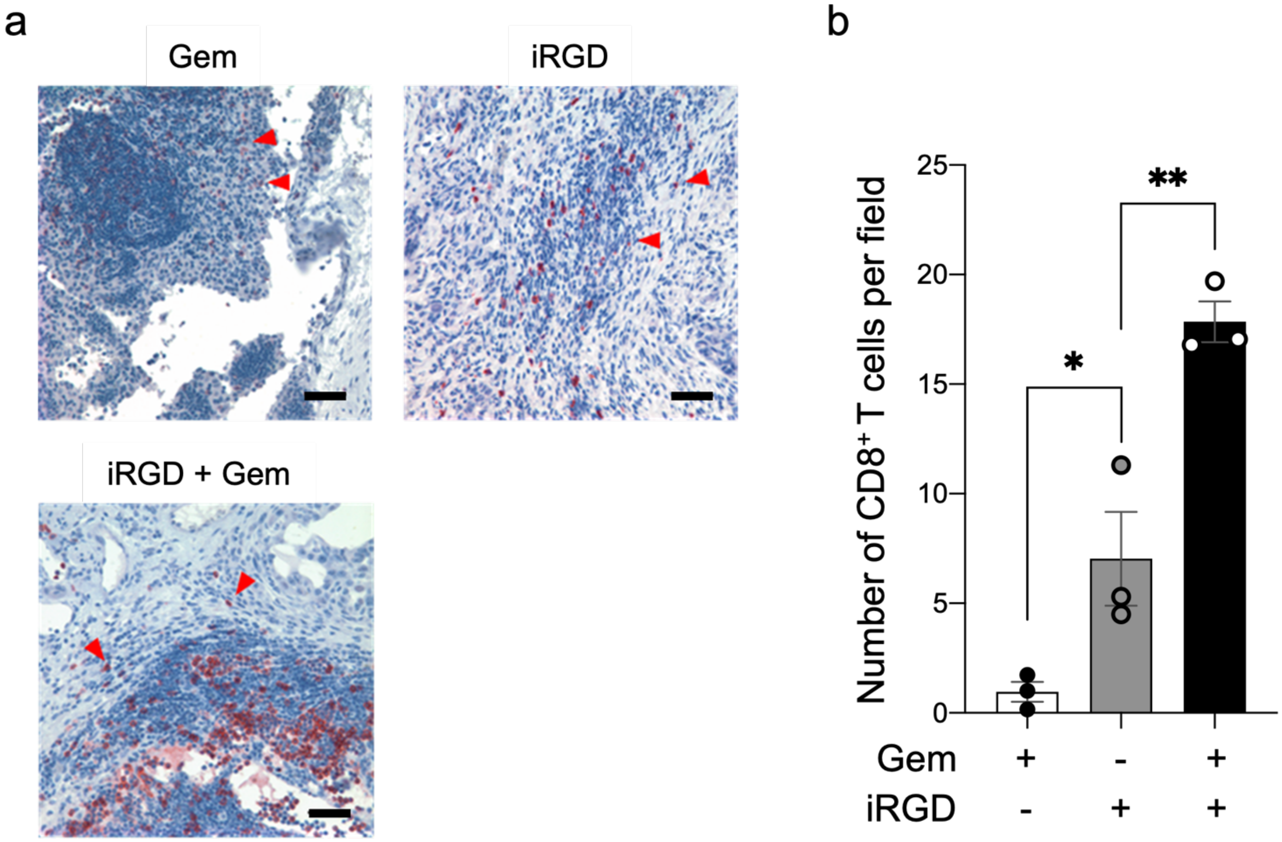


**Fig. S1 - Long-term iRGD therapy increases CD8$^+$ T cells in the PDAC tissue in transgenic KPC mice.** Primary PDAC tumors were collected from transgenic KPC mice that received long-term treatment with Gem, iRGD, or iRGD + Gem as part of a previously published survival study[25]. The primary PDAC tumors were immunohistochemically analyzed for the presence of CD8$^+$ T cells. **a** Representative images of CD8$^+$ T cells in the PDAC tissue (brown, arrowheads show examples). Scale bars, 50 µm. **b** A bar diagram showing the number of CD8$^+$ T cells per high-power field. n = 3 per arm. Statistical analysis, one-way ANOVA; $p = 0.0468$ (Gem vs iRGD), $p = 0.0034$ (iRGD vs Gem + iRGD). Error bars, mean ± standard error; $*p < 0.05$; $**p < 0.01$.

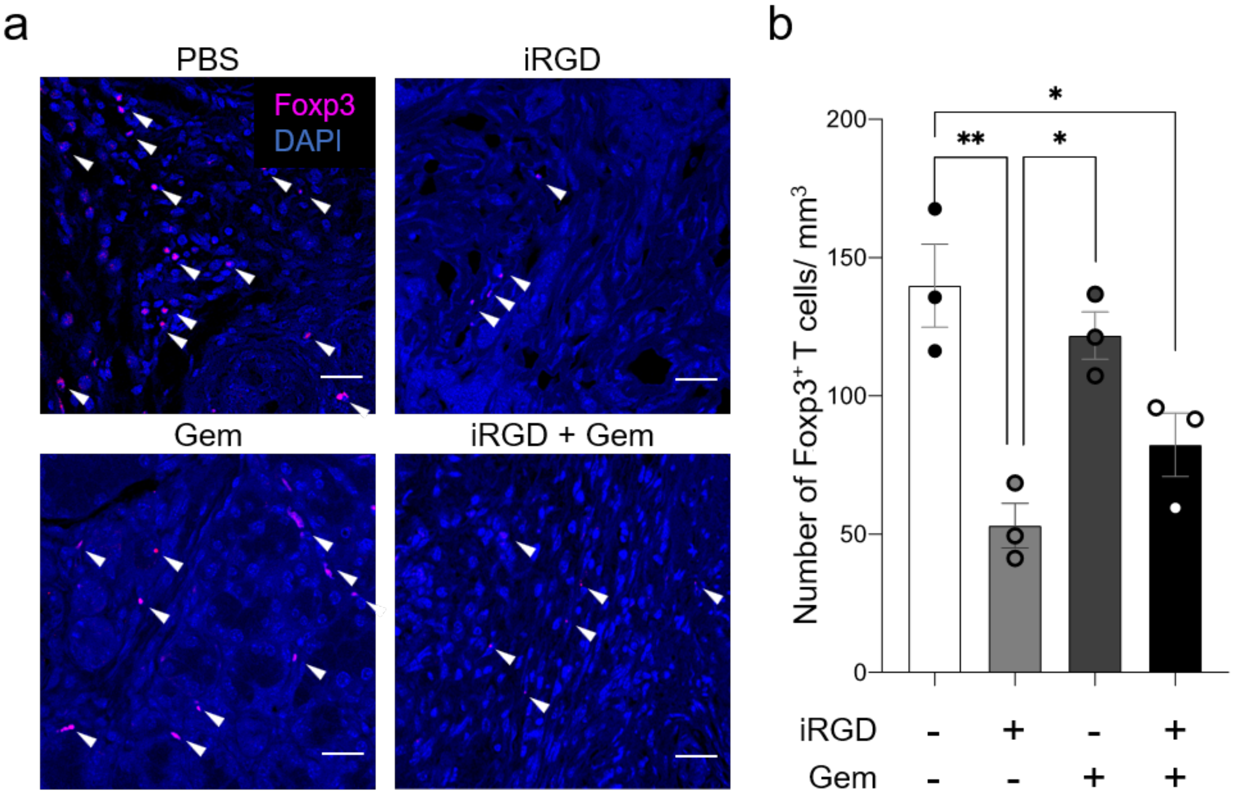


**Fig. S2 - Long-term iRGD therapy decreases Foxp3$^+$ cells in the PDAC in transgenic KPC mice.** Primary PDAC tumors were collected from transgenic KPC mice that received long-term treatment with or without Gem, iRGD, or iRGD + Gem as part of a previously published survival study[25]. The presence of Foxp3$^+$ cells in the primary PDAC tumors was analyzed by immunofluorescence. **a** Representative images of Foxp3$^+$ cells (red) in the PDAC tissue. Blue, DAPI; scale bars, 25 µm. **b** A bar diagram showing the number of Foxp3$^+$ cells per high-power field in the primary PDAC tumors. n = 3 per arm. Statistical analysis, one-way ANOVA; $p = 0.0025$ (PBS vs iRGD), $p = 0.0101$ (iRGD vs Gem), $p = 0.0261$ (PBS vs Gem + iRGD). Error bars, mean ± standard error; $^*p < 0.05$; $^{**}p < 0.01$.

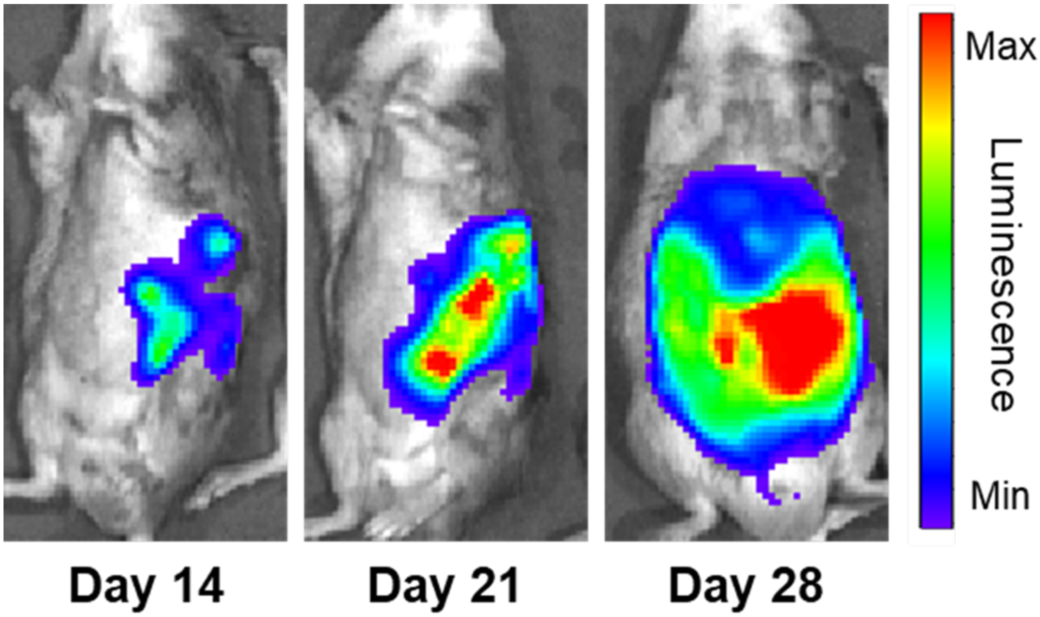


**Fig. S3 - Rapid growth of orthotopic KPC-derived PDAC tumors in B6129SF1/J hybrid mice.** Representative images showing longitudinal growth of an orthotopic luciferase$^{+}$ KPC-derived PDAC tumor in a mouse with a matching genetic background analyzed by bioluminescence imaging.

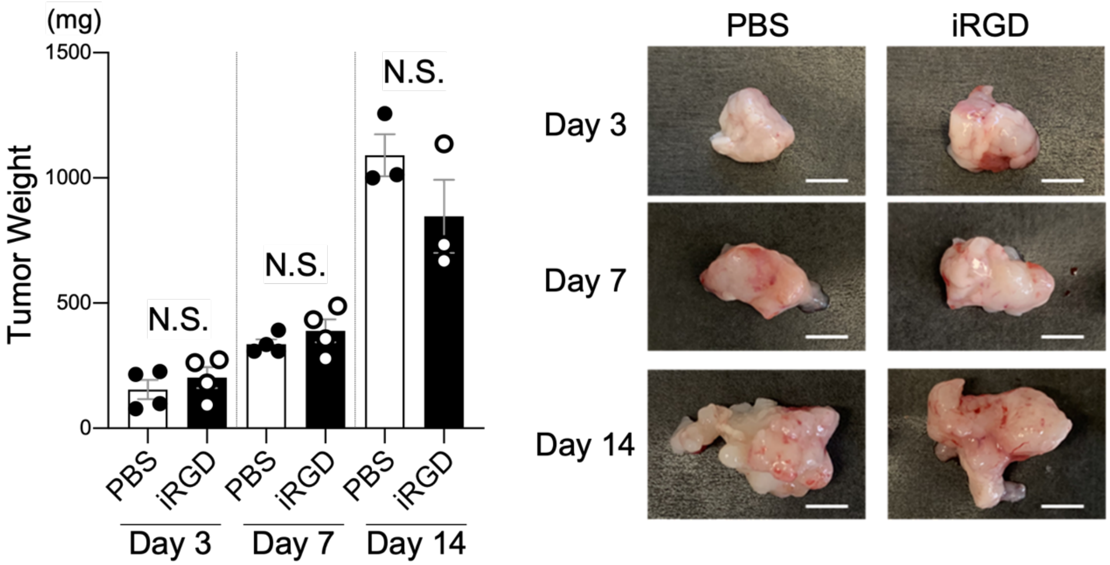


**Fig. S4 - iRGD monotherapy does not affect PDAC growth.** C57B6129SF1/J hybrid mice bearing orthotopic KPC-derived PDAC tumors were treated with PBS or iRGD 3 times a week for 2 weeks. The tumors were collected and weighed 3, 7 and 14 days after the treatment was started. The results are summarized in the bar diagram. n = 4 (day 3 and day 7); n = 3 (day 14). Statistical analysis, Mann-Whitney U test (day 3 and day 7) or Welch's *t* test (day 14); $p = 0.4857$ (day 3), $p = 0.4857$ (day 7), $p = 0.2389$ (day 14). Error bars, mean ± standard error; N.S., not significant. Representative images of the tumors are shown to the right. Scale bars, 5 mm.

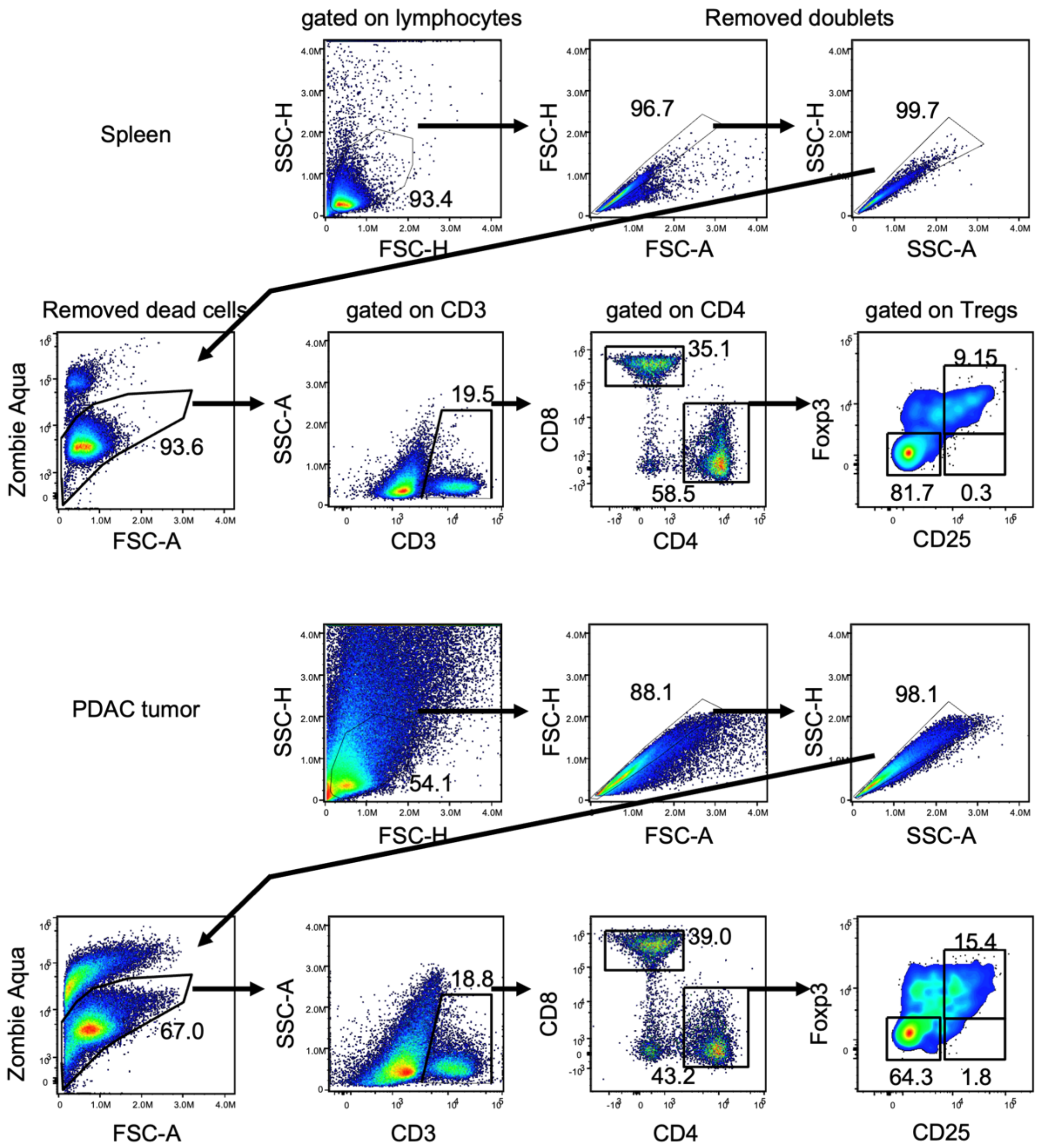


**Fig. S5 – Gating strategy for flow cytometric analysis of splenic and tumor-resident T cells in orthotopic KPC-derived PDAC mice.** Immune cells isolated from the spleens and tumors of C57B6129SF1/J hybrid mice bearing KPC-derived orthotopic PDAC were subjected to flow cytometry. Lymphocytes were gated, and doublets and dead cells were excluded. CD3$^+$ T cells were separated by CD4 and CD8 positivity, and the CD4$^+$ T cells were expanded using CD25 and Foxp3. As CD4$^+$ CD25$^+$ T cells were nearly universally positive for Foxp3, CD4$^+$ CD25$^+$ T cells were defined as Tregs for flow cytometry studies of tumor-resident T cells. Upper panels, splenocytes; lower panels, tumor-resident T cells.

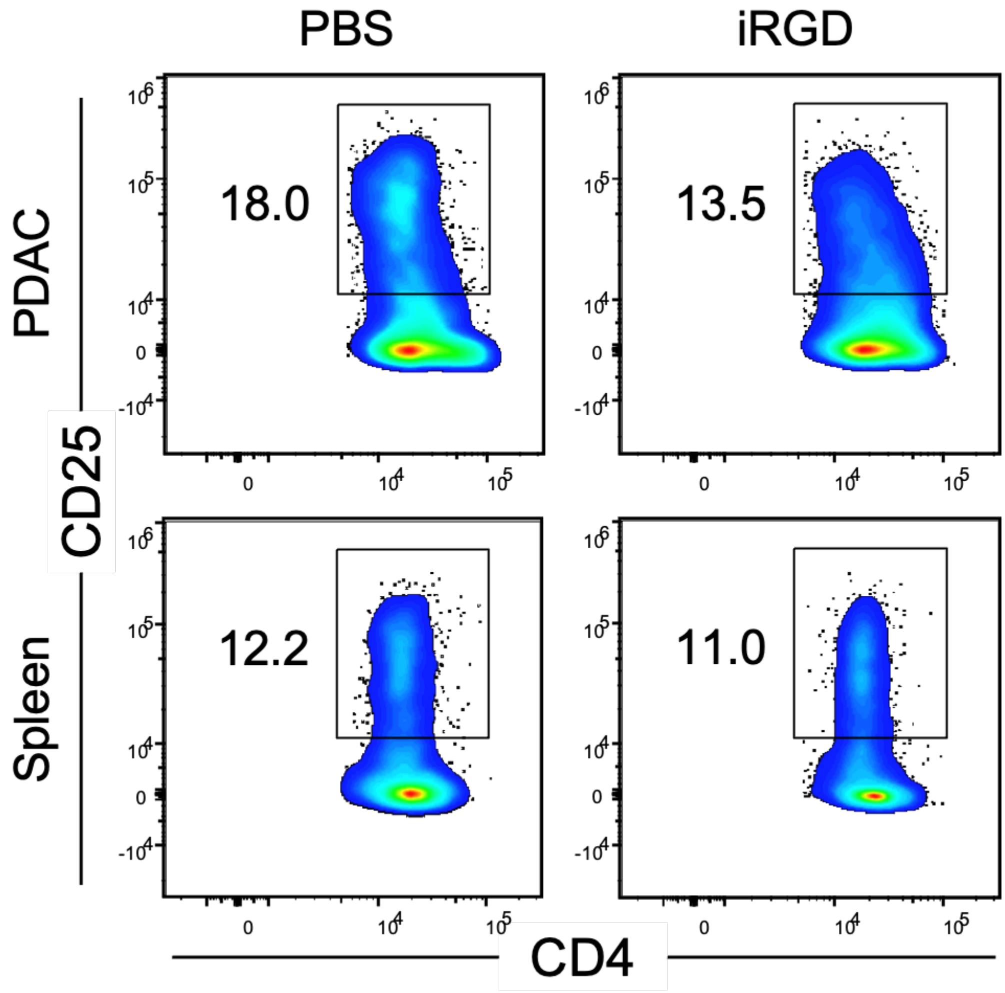


**Fig. S6 – Changes in the proportion of Tregs in the PDAC and spleen of KPC-derived PDAC mice upon iRGD monotherapy.** C57B6129SF1/J hybrid mice bearing KPC-derived orthotopic PDAC tumors were intravenously treated with PBS or iRGD 3 times a week for 2 weeks. Tumors and spleens were collected 3, 7 and 14 days after the treatment was started. The proportion of $CD4^+$ $CD25^+$ Tregs among $CD4^+$ T cells in the tumor and the spleen was analyzed by flow cytometry. Representative dot plots from day 14 are shown. The results are summarized in Figs. 1a and 1c.

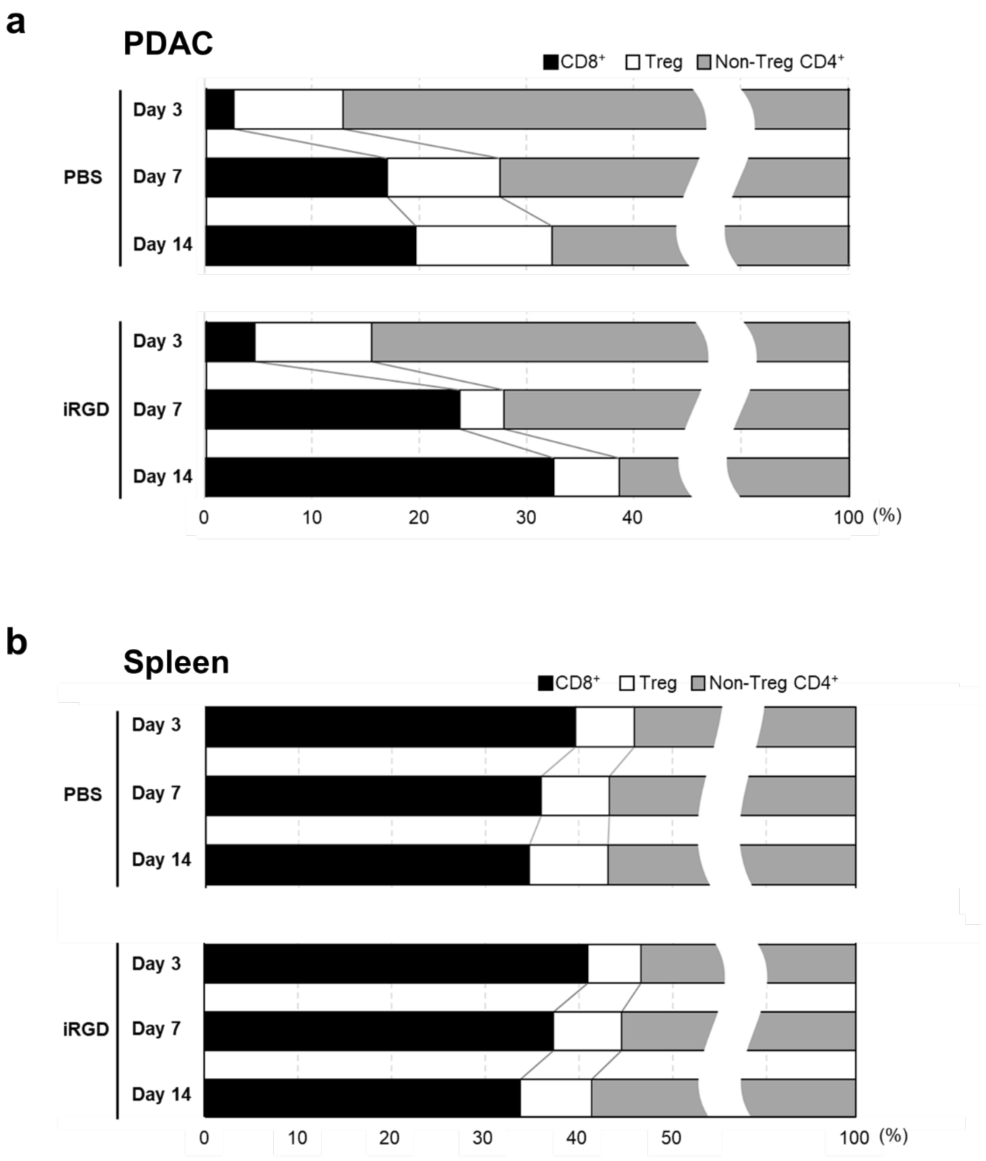


**Fig. S7 - The effect of iRGD therapy on the proportion of $CD8^+$ T cells, Tregs, and non-Treg $CD4^+$ T cells in the PDAC tissue and the spleen.** C57B6129SF1/J hybrid mice bearing orthotopic KPC-derived PDAC tumors were treated with PBS or iRGD 3 times a week for 2 weeks. The tumors and spleens were collected 3, 7 and 14 days after the treatment was started. The proportion of $CD8^+$ T cells, $CD4^+$ $CD25^+$ T cells (Tregs), and $CD4^+$ $CD25^{neg}$ T cells (non-Tregs) in the tumors (**a**) and the spleens (**b**) was analyzed by flow cytometry. n = 3 per arm.

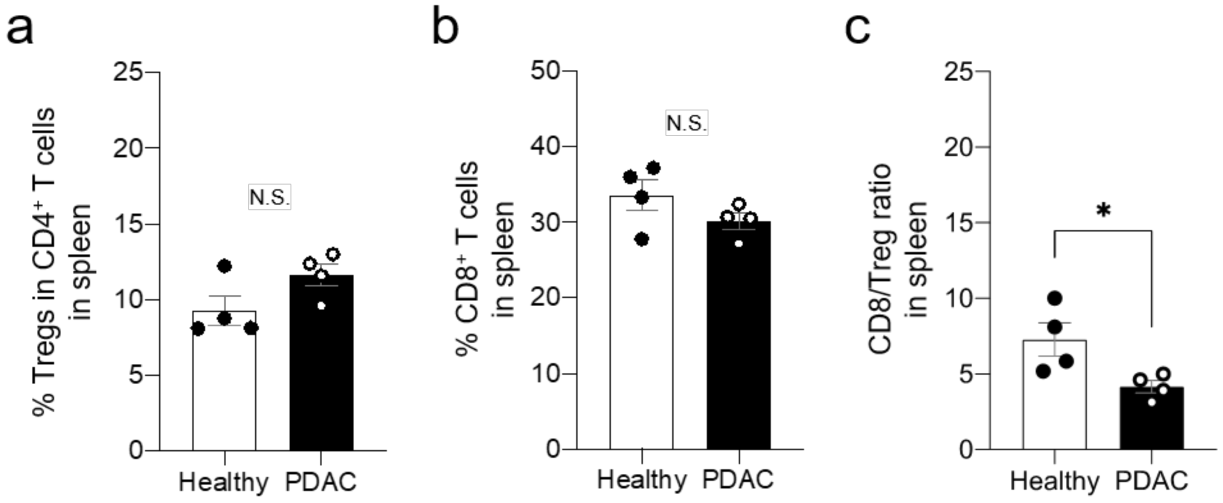


**Fig. S8 – The proportion of Tregs in $CD4^+$ T cells and the CD8/Treg ratio in the spleen of healthy mice and PDAC mice.** Spleens from healthy C57B6129SF1/J hybrid mice and C57B6129SF1/J mice bearing 22 day-old orthotopic KPC-derived PDAC tumors were collected. Splenocyte were isolated and subjected to flow cytometry. **a** The proportion of $CD4^+$ $CD25^+$ Tregs among $CD4^+$ T cells. **b** The proportion of $CD8^+$ T cells. **c** The CD8/Treg ratio was calculated based on the values from **a** and **b**. n = 4 per group. Statistical analysis, Mann-Whitney U test; $p = 0.1143$ (**a**), $p = 0.2000$ (**b**), $p = 0.0286$ (**c**). Error bars, mean ± standard error; *$p < 0.05$; N.S., not significant.

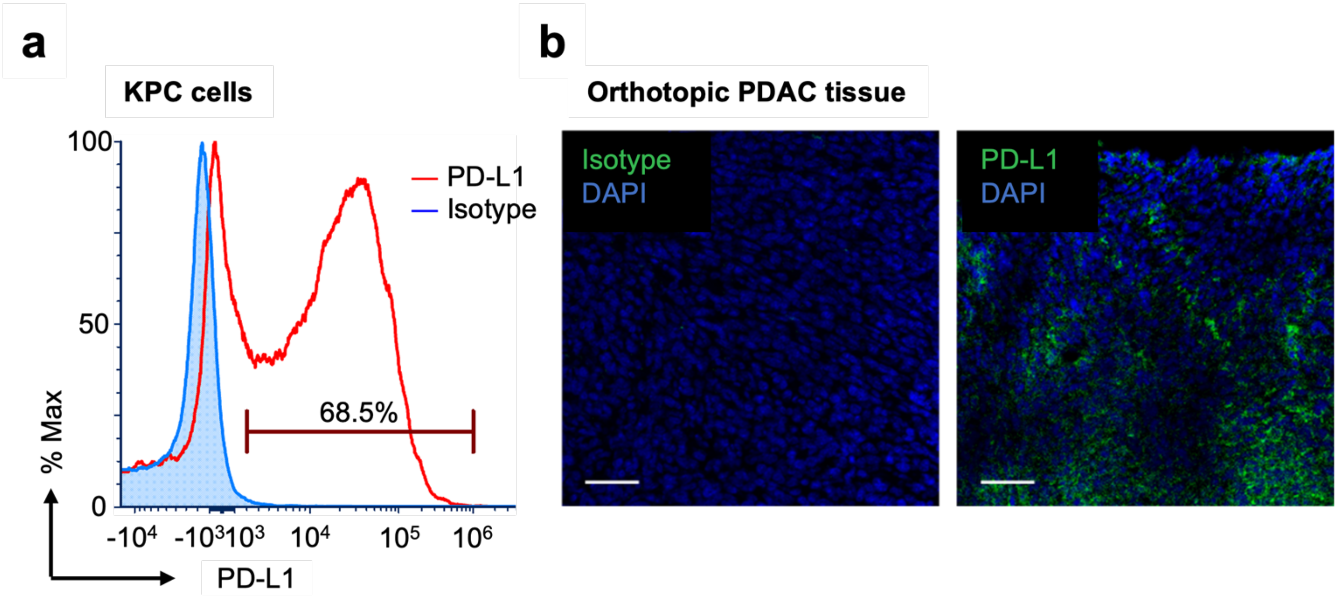


**Fig. S9 - KPC-derived PDAC cells express high levels of PD-L1. a** PD-L1 expression on KPC-derived PDAC cells analyzed by flow cytometry. Blue, isotype control; red, anti-PD-L1 Ab. **b** Representative immunofluorescence images showing the expression of PD-L1 (green) in orthotopic KPC-derived PDAC tissue that was harvested 22 days after tumor cell inoculation. Blue, DAPI. Scale bars, 50 μm.

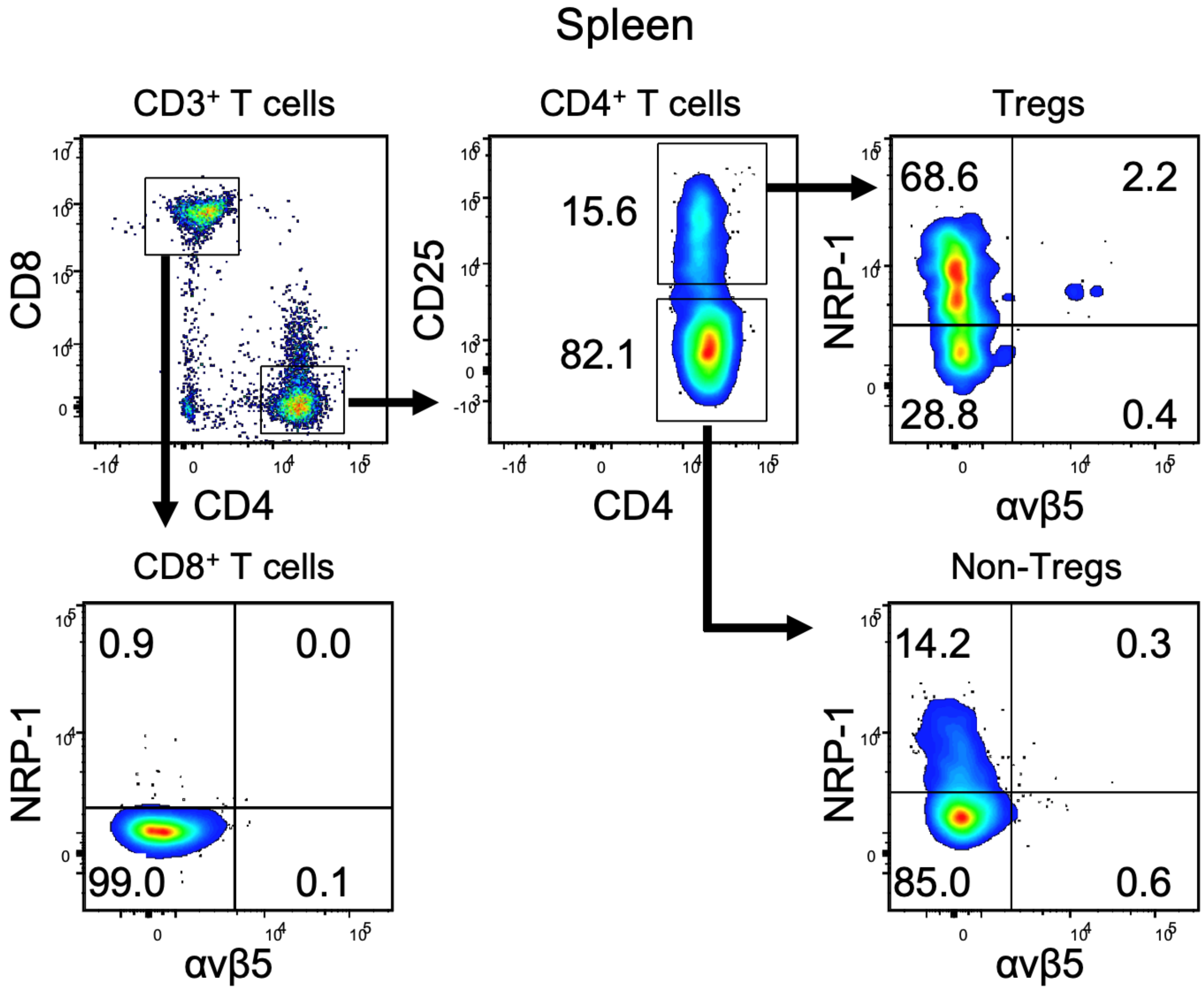


**Fig. S10 – Representative flow cytometry analysis of αvβ5 integrin and NRP-1 expression on splenic T cells.** A representative flow cytometry analysis showing the proportion of $CD8^{+}$ T cells, $CD4^{+}$ $CD25^{neg}$ T cells (non-Tregs), $CD4^{+}$ $CD25^{+}$ Tregs that are positive for αvβ5 integrin, NRP-1, or both in the spleen of mice bearing KPC-derived orthotopic PDAC.

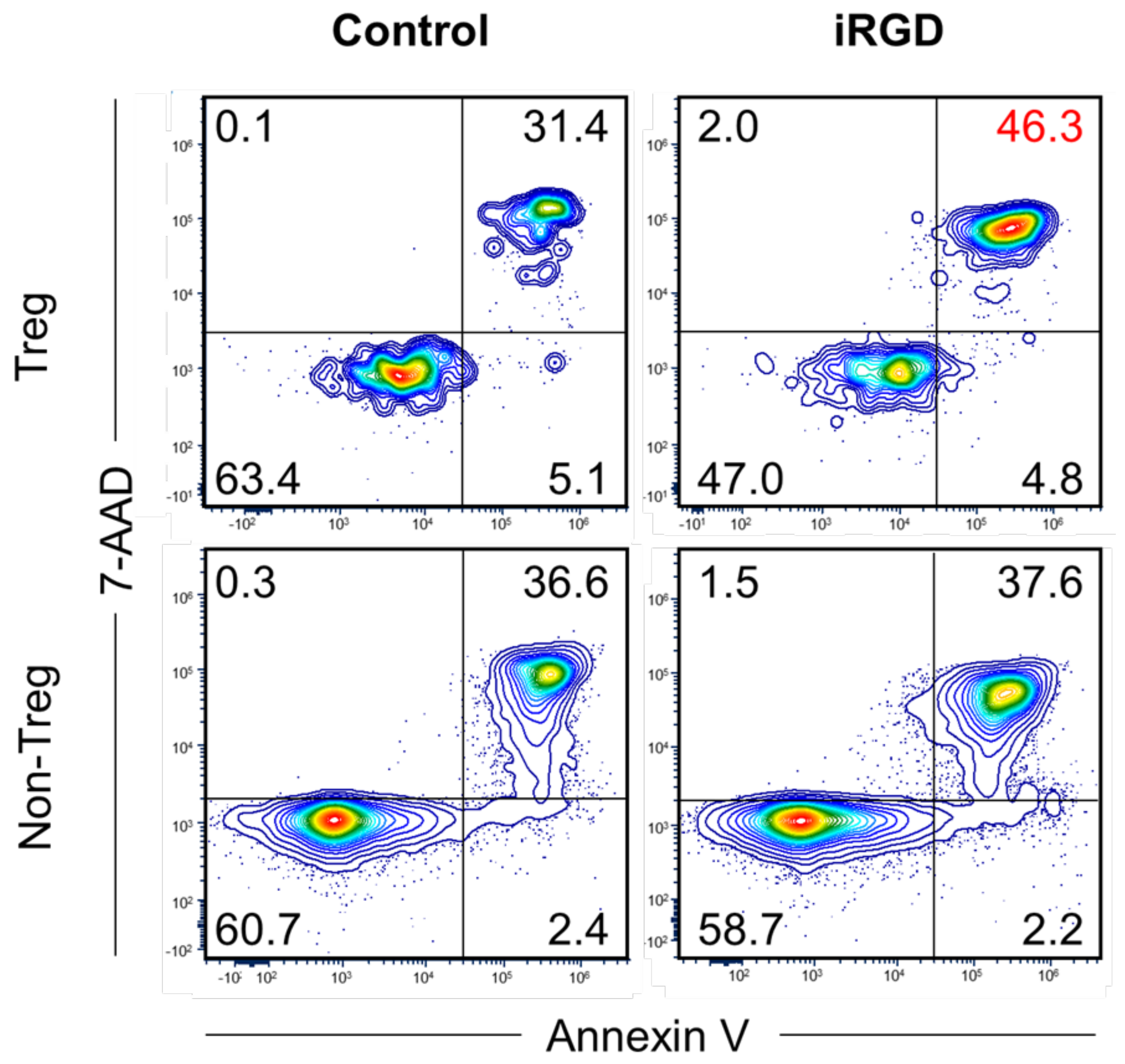


**Fig. S11 – iRGD induces apoptosis of Tregs in the presence of KPC-derived PDAC cells.** $CD4^+$ T cells isolated from the spleens of healthy C57B6129SF1/J hybrid mice were expanded for 3 days *in vitro* in the presence of KPC-derived PDAC cells and iRGD. Apoptosis of $CD4^+$ $CD25^+$ Tregs and $CD4^+$ $CD25^{neg}$ T cells (non-Tregs) was quantified by measuring annexin V and 7-AAD double positive cells by flow cytometry. Representative dot plots are shown. The results are summarized in Fig. 3e.

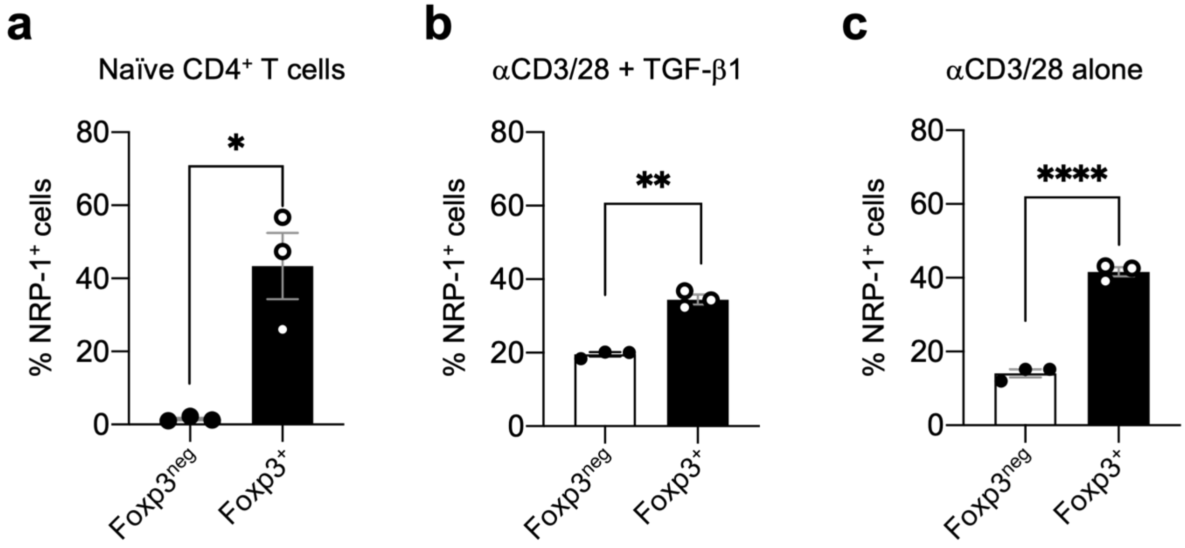


**Fig. S12 – NRP-1 expression on naïve CD4$^+$ T cells before and after TCR stimulation with or without TGF-β1.** A pool of CD4$^+$ T cells enriched for naïve CD4$^+$ T cells were treated with anti-CD3/CD28 Abs in the presence or absence of TGF-β1 and subjected to flow cytometry as described in Fig. 5. **a** NRP-1 expression on naïve CD4$^+$ Foxp3$^{neg}$ T cells and CD4$^+$ Foxp3$^+$ T cells prior to the treatment. **b, c** NRP-1 expression on CD4$^+$ Foxp3$^{neg}$ T cells and CD4$^+$ Foxp3$^+$ T cells after treatment with anti-CD3/CD28 Abs and TGF-β1 (**b**) or with anti-CD3/CD28 Abs alone (**c**). n = 3 per study. Statistical analysis, Welch's *t* test; $p = 0.0435$ (**a**), $p = 0.0027$ (**b**), $p < 0.0001$ (**c**). Error bars, mean ± standard error; *$p < 0.05$; **$p < 0.01$; ****$p < 0.0001$.

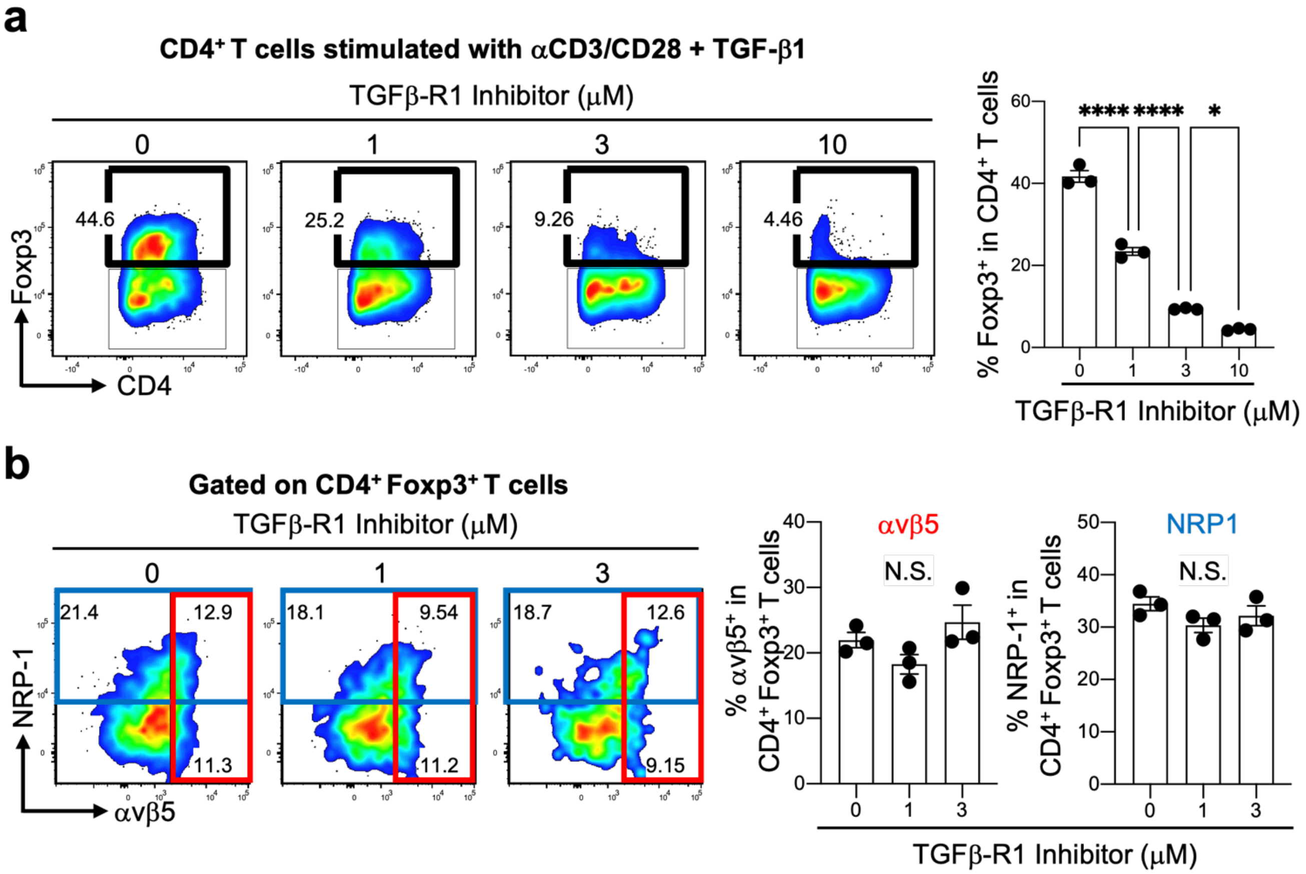


**Fig. S13 – TGF-βR1 inhibitor does not affect the expression of αvβ5 integrin and NRP-1 on iTregs.** Naïve $CD4^+$ T cells isolated from the spleens of healthy mice were expanded in the presence of anti-CD3/CD28 Abs, exogenous TGF-β1, and increasing doses of a TGF-βR1 inhibitor (LY2157299). **a** Flow cytometric analysis showing the proportion of $CD4^+$ $Foxp3^+$ T cells (iTregs) after the expansion. The bar diagram summarizes the results from 3 independent experiments. Statistical analysis, One-way ANOVA; $p < 0.0001$ (0 vs 1), $p < 0.0001$ (1 vs 3), $p = 0.0154$ (3 vs 10). **b** Expression of αvβ5 integrin (red boxes) and NRP-1 (blue boxes) on the iTregs gated in (**a**). The bar diagram summarizes the results from 3 independent experiments. Statistical analysis, one-way ANOVA; $p = 0.3996$ (αvβ5, 0 vs 1), $p = 0.5868$ (αvβ5, 0 vs 3), $p = 0.2265$ (NRP-1, 0 vs 1), $p = 0.5751$ (NRP-1, 0 vs 3). Error bars, mean ± standard error; *$p < 0.05$; ****$p < 0.0001$; N.S., not significant.

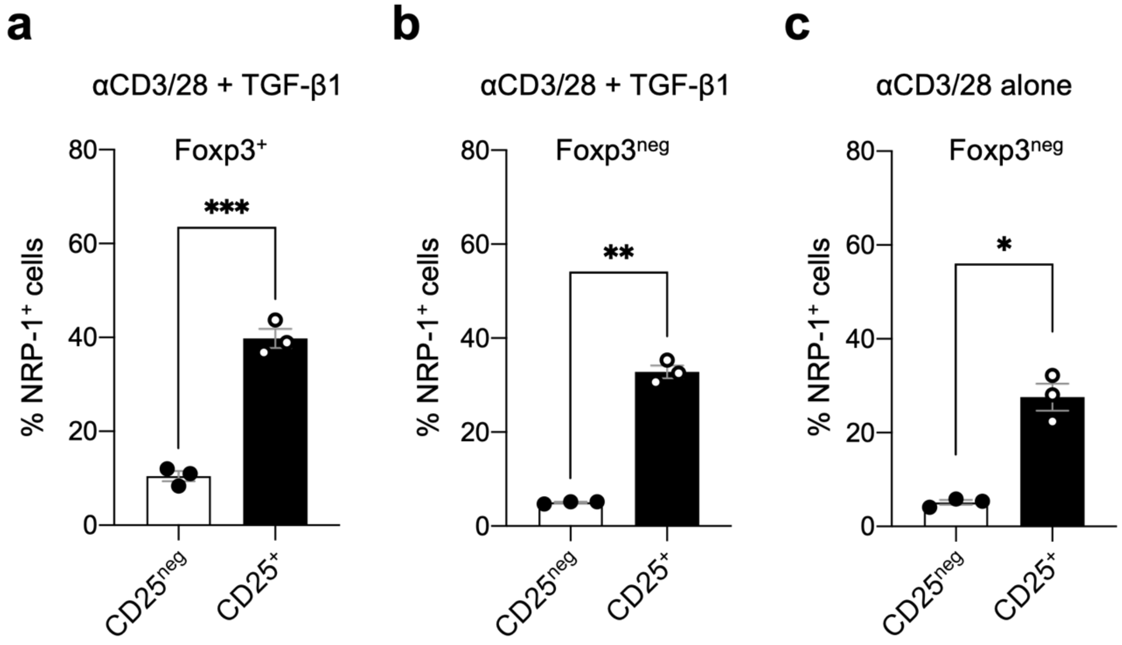


**Fig. S14 – Relevance of NRP-1 and CD25 expression profiles in naïve CD4$^+$ T cells that received TCR stimulation with or without TGF-β1.** A pool of CD4$^+$ T cells enriched for naïve CD4$^+$ T cells were treated with anti-CD3/CD28 Abs in the presence (**a, b**) or absence (**c**) of TGF-β1 and subjected to flow cytometry as described in Fig. 6. The bar diagrams show the proportion of NRP-1$^+$ cells among CD25-positive and negative populations in CD4$^+$ Foxp3$^+$ T cells (**a**) and CD4$^+$ Foxp3$^{neg}$ T cells (**b, c**) after the treatment. n = 3 per study. Statistical analysis, two-tailed unpaired Welch's *t* test; $p = 0.001$ (**a**), $p = 0.0021$ (**b**), $p = 0.0135$ (**c**). Error bars, mean ± standard error; *$p < 0.05$; **$p < 0.01$.

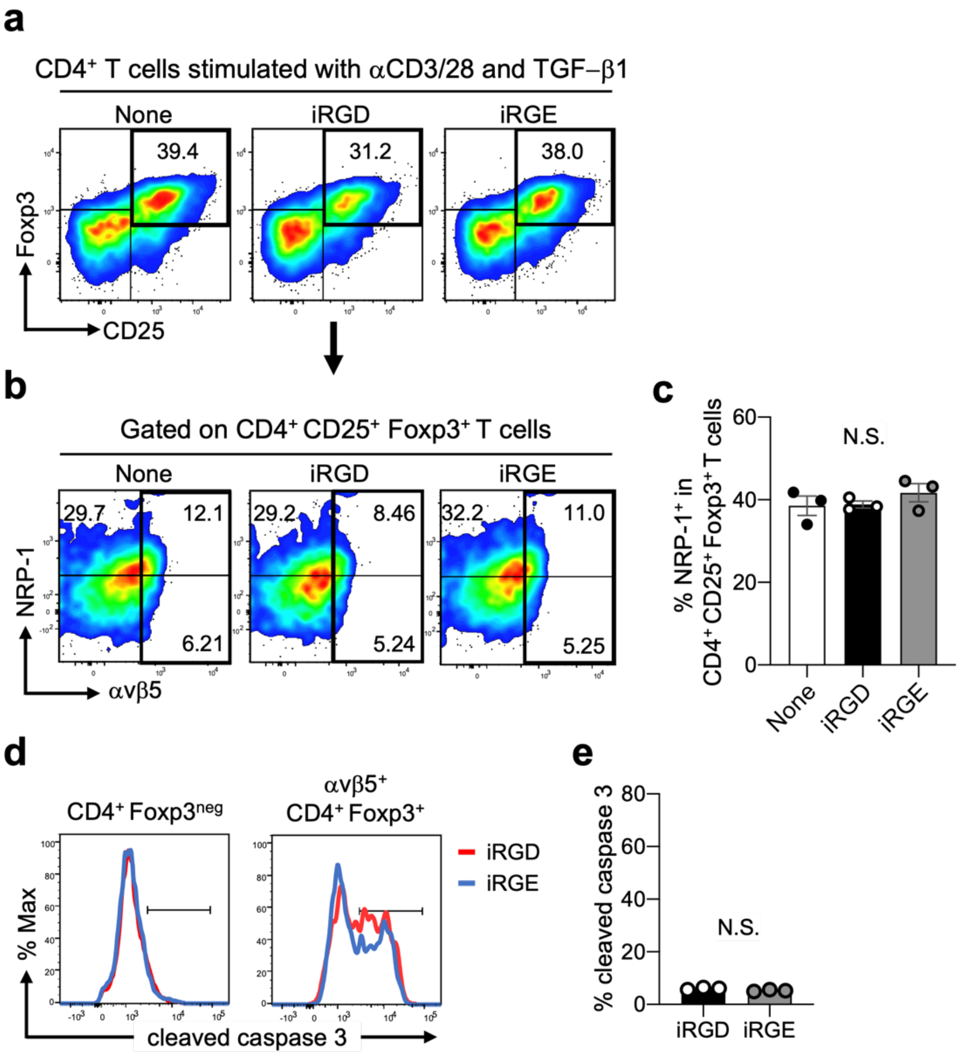


**Fig. S15 - iRGD reduces $CD4^+$ $CD25^+$ $Foxp^+$ iTregs in an RGD-dependent manner.** Naïve $CD4^+$ T cells were stimulated with anti-CD3/CD28 Abs and TGF-β1 in the absence or presence of the iRGD or iRGE peptide. **a-c** The resulting cells were subjected to flow cytometry to analyze the expression of CD25 and Foxp3 in the $CD4^+$ T cells (**a**) and the expression of the αvβ5 integrin and NRP-1 on $CD4^+$ $CD25^+$ $Foxp3^+$ iTregs (**b**) The bar diagram in (**c**) summarizes the proportion of $NRP\text{-}1^+$ cells among the iTregs. Statistical analysis, one-way ANOVA; $p = 0.8221$. n = 3 per group. **d**, **e** The proportion of apoptotic αvβ5 $integrin^+$ iTregs (**d**) and $CD4^+$ $Foxp3^{neg}$ T cells (**e**) was quantified by measuring cleaved caspase 3 using flow cytometry. Statistical analysis, Welch's *t* test; $p = 0.0705$. n=3. The dot plots and histograms show representative data from the 3 independent experiments. Error bars, mean ± standard error; N.S., not significant.

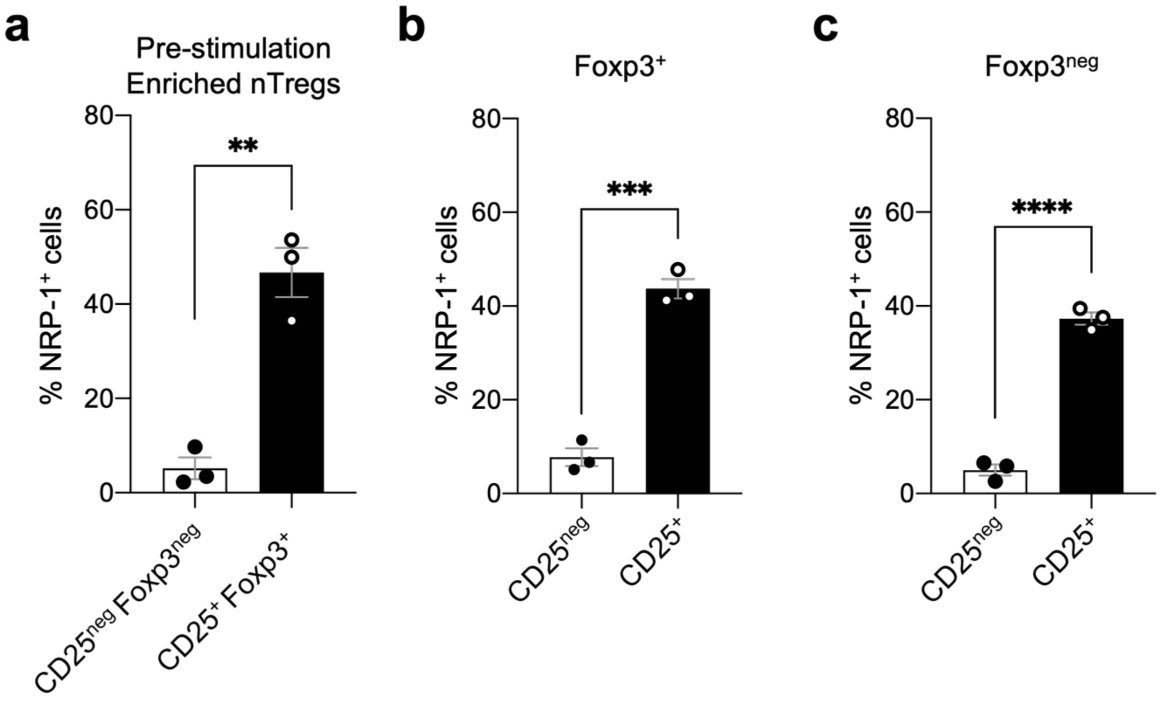


**Fig. S16 – NRP-1 expression on nTregs before and after TCR stimulation.**

A pool of CD4$^+$ T cells enriched for CD4$^+$ CD25$^+$ Foxp3$^+$ nTregs were treated with anti-CD3/CD28 Abs and subjected to flow cytometry as described in Fig. 7. **a** NRP-1 expression on the nTregs and CD4$^+$ CD25$^{neg}$ Foxp3$^{neg}$ T cells prior to the treatment. **b, c** NRP-1 expression on CD25-positive and negative populations in CD4$^+$ Foxp3$^+$ T cells (**b**) and CD4$^+$ Foxp3$^{neg}$ T cells (**c**) analyzed after the treatment. n = 3 per study. Statistical analysis, Welch's *t* test; $p = 0.007$ (**a**), $p = 0.0002$ (**b**), $p < 0.0001$ (**c**). Error bars, mean ± standard error; **$p < 0.01$; ***$p < 0.001$; ****$p < 0.0001$.
